# Evidence of genetic isolation and differentiation among historically fragmented British populations of common juniper, *Juniperus communis* L.

**DOI:** 10.1101/2024.09.26.615127

**Authors:** J. Baker, J. Cottrell, R. Ennos, A. Perry, S. A’Hara, S. Green, S. Cavers

## Abstract

Habitat fragmentation and populations isolation pose a threat to the genetic diversity and adaptability of many species. The common juniper, Juniperus communis L., a keystone species for juniper scrub habitat and one of only three conifers that are native to the UK, has been in decline for more than a century in the UK and across its European range. Remnant UK juniper populations are now highly fragmented and often small, which has raised concerns for their resilience, especially in the face of climate change and the introduction of novel pathogens, such as Phytophthora austrocedri. This work presents a baseline genetic survey of native UK juniper populations and compares patterns of diversity between populations and among three population centres in southern England, the Lake District, and Scotland using both Single Nucleotide Polymorphism (SNP) and Simple Sequence Repeat (SSR) genetic markers. The aim was to evaluate the standing genetic diversity of native juniper stands, the impacts of habitat fragmentation, and to determine whether juniper populations are genetically isolated from one another. We found that juniper populations, while not completely isolated from one another, face substantial barriers to gene flow, especially between the three population centres. These centres also show different patterns of genetic diversity, indicating varying levels of internal gene flow and inbreeding. Our findings can form a baseline from which to monitor the effectiveness of conservation activities, prioritize populations of concern, and guide genetic rescue efforts.

## Introduction

**Figure 1:**
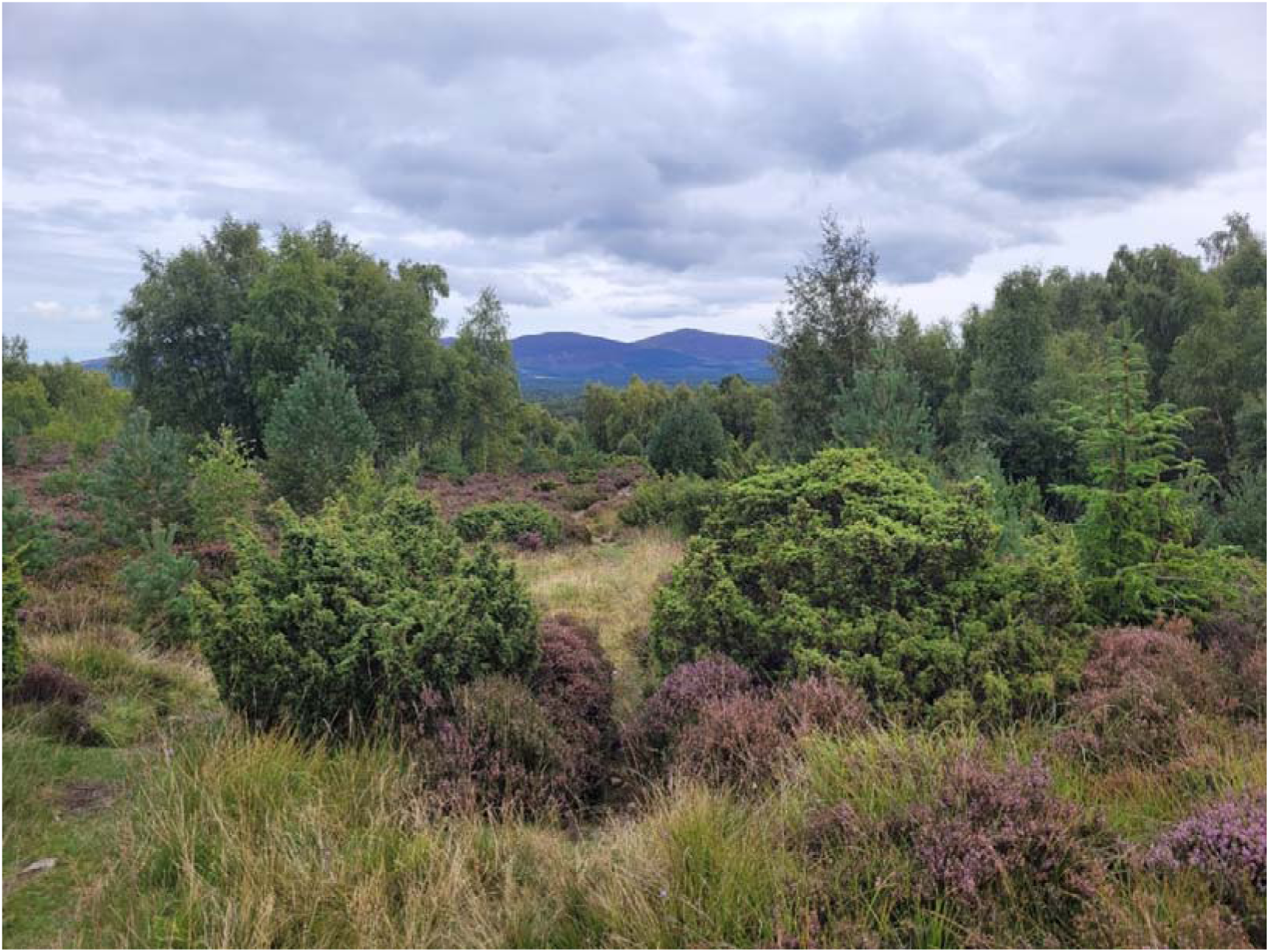
Cover photo taken by Baker, J.

Habitat fragmentation represents a threat to the genetic diversity of tree species as it can lead to a decline in gene flow between populations and a resultant increase in genetic drift and inbreeding in remnant populations (Aguilar et al., 2008; Dobeš et al., 2017; Young et al., 1996). The potential reduction of genetic diversity in fragmented populations may in turn impair the adaptive potential of populations (Cavers & Cottrell, 2015; Ennos, 2015).

Generally, features of a population that facilitate the production of new genotypes, such as larger population size, inter-population gene flow, and abundant natural regeneration, all maintain or increase the adaptive potential of that population. Therefore, conservation with the explicit goal of maintaining or increasing the genetic diversity of species and populations by promoting gene flow and natural selection, often called dynamic conservation, has become a recognized method with which to create more resilient forest populations (Cavers & Cottrell, 2015; Fady et al., 2016; Finger et al., 2022; Hubert & Cottrell, 2014; Lefèvre et al., 2013). The effects of habitat fragmentation are not ubiquitous and are determined by mating systems, life-history traits and the demographics of the meta-population before the fragmentation occurred (Aguilar et al., 2008; Lowe et al., 2015). It is therefore beneficial to understand the impacts of fragmentation on a species through the evaluation of both the direct impacts (e.g. inbreeding, population structure) *via* population genetics, and indirect impacts (e.g. loss of phenotypic diversity or distribution of locally adaptive traits) *via* quantitative genetics (Cavender-Bares & Ramírez-Valiente, 2017; De Kort et al., 2013; de Villemereuil et al., 2016; Ennos et al., 1998). Although temperate tree species, such as Scots pine, have generally been found to have very high levels of gene flow between populations (Rodriguez, 2019; Salmela, 2011; Salmela et al., 2013), some species, such as yew, seem to be more sensitive to fragmentation (Chybicki et al., 2024). Here, we use a novel set of Sequenom Single Nucleotide Polymorphisms (SNPs) and a newly developed panel of Simple Sequence Repeats (SSRs) to quantify the genetic diversity and infer the effects of habitat fragmentation on the keystone species the common juniper, *Juniperus communis* L.

The common juniper has the widest global distribution of any conifer species, with a circumpolar range that extends from northern tundra in Russia and Canada as far south as the Mediterranean in Europe and the Central Rockies in North America (Thomas et al., 2007). The species is morphologically variable, and can grow as upright mid-story trees, sprawling shrubs, or ground-hugging stems (Carrer et al., 2019; Klimko et al., 2007; Knyazeva & Hantemirova, 2020). It is dioecious, wind-pollinated, and its seeds are primarily dispersed by birds (Adams & Thornburg, 2010; García, 2001; Surso, 2018; Thomas et al., 2007). Juniper trees are a keystone species for many of the communities in which they occur, are particularly important as habitats for lichens and bryophytes, and provide abundant seasonal forage for animals. Furthermore, they can aid the recruitment of other tree species by acting as “nursery trees,” protecting young tree seedlings when they are particularly vulnerable to predation.

Juniper is one of only three conifers native to the UK, and its extracts have and continue to be used, for both medical and culinary purposes, since at least 1550 B.C.E. (Al-Snafi, 2018).

Although juniper’s considerable phenotypic variability and dispersal strategies (Hall, 1990; Knyazeva & Hantemirova, 2020; Thomas et al., 2007) might suggest a highly adaptable and resilient species, populations have been declining for at least the past century in both the UK and elsewhere in Europe. Consequently, juniper is listed as a priority species under the UK’s Action Plan for Biodiversity (McBride, 2005) and many juniper communities are listed as Special Areas of Conservation under the EU’s Habitats Directive. In the UK, remnant juniper populations are generally small and some, such as those in Southern England, are failing to regenerate naturally. Changes in land use, particularly grazing and the absence of regular natural disturbances (De Frenne et al., 2020; McBride, 2005; Thomas et al., 2007), are considered the primary reasons for the lack of seedling recruitment, but there are many other factors that may contribute, including increasing temperatures (R. Gruwez et al., 2016; Verheyen et al., 2009) and changing soil nutrient compositions (Gruwez et al., 2014; Robert Gruwez et al., 2016; Pers-Kamczyc et al., 2022; Pers-Kamczyc et al., 2020; Verheyen et al., 2009). The lack of natural regeneration is resulting in populations that are both shrinking and aging with more male-biased sex ratios, which is especially concerning given that reproductive success may decrease as plants age (Garcı a et al., 1999; Ward, 1982, 2007).

Previous genetic surveys of junipers have typically been restricted to relatively small geographic areas when compared to juniper’s global range, and often differ in the genetic markers used, making direct comparisons difficult. However, Knyazeva and Hantemirova (2020) studied the population and quantitative genetics of juniper across its entire range in Russia. Our study is the first to include samples from both Scotland and England and includes all of the subspecies that occur on the British Isles: *J. communis* spp. *communis, J. communis* spp. *nana*, and *J. communis* spp. *hemisphaerica* (hereafter abbreviated as *J. communis, Nana* and *Hemi*, respectively). The three subspecies are primarily distinguished by leaf morphology (Stace, 2019) although *J. communis* and *Nana* may also be distinguished by their cone anatomy (Sullivan, 2001). The three subspecies also differ in their ranges, with *Nana* being restricted to the west coast of Scotland and Wales (Sullivan, 2003; Thomas et al., 2007) and *Hemi* being restricted to a single population in Cornwall (Stace, 2019; Thomas et al., 2007). Although the genetic status of the subspecies is not clear, Sullivan (2001) found evidence that the prostrate growth habit of *Nana* was a genetic adaptation that was retained in a common garden trial, whereas prostrate *J. communis* cuttings demonstrated some phenotypic plasticity in their growth habits when grown in a common garden trial. Sullivan (2001) did not, however, find support for their genetic distinction based on RAPD markers. Similarly, the genetic status of *Hemi* is unclear, and it is often regarded as an intermediate between the other two subspecies (Thomas et al., 2007), however there is some evidence of its genetic distinction from the other subspecies based on AFLP markers (Merwe et al., 2000).

The goal of this work was to provide a genetic survey that allows for the comparison of the larger Scottish populations with the smaller and more fragmented ones in the Lake District and southern England. This work uses both SNP and SSR genetic markers, as they are complementary approaches with different strengths and weaknesses (García et al., 2018). Quantifying the patterns of genetic diversity in these neutral genetic markers can inform researchers and conservationists about the population-scale dynamics of gene flow, the effects of habitat fragmentation and the development of effective management strategies. Our aim was to provide guidance to conservation managers in Britian and to target the selection of British some juniper populations as Gene Conservation Units (GCUs) under the European Forest Genetic Resources Programme (EUFORGEN).

## Methods

### Population locations, material collection and DNA extraction

Sixteen populations of *J. communis* and one each of *Nana* and *Hemi* were sampled from sites in England and Scotland in this study (Table 1 and Figure 2). Although the UK distribution of juniper includes populations in Wales and Ireland, these were not included in this study. Needle samples were collected in 2019 and stored in −20 LC prior to processing.

**Figure 2:**
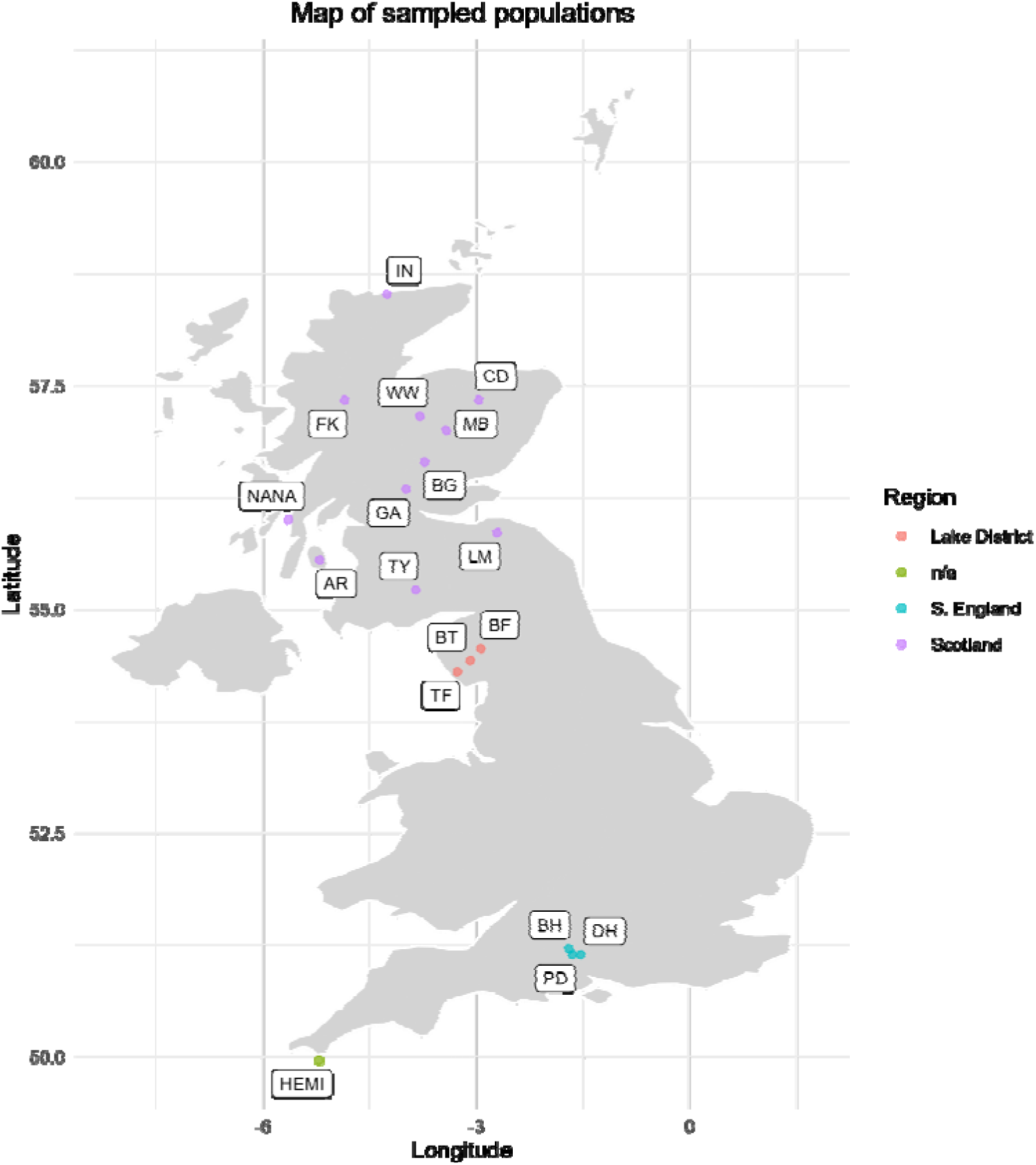
Map displaying locations of sampled populations, demarked with coloured circles corresponding to the region and with population name abbreviations listed in Table 1.

**Table 1:** List of populations that were included in genetic analyses, including the abbreviations that are used in visuals, coordinates and the number (N) of individuals in both SNP and SSR datasets.

| Population | Abbreviation | Region | Lat (°) | Long (°) | N<br>for<br>SNP | N for<br>SSR |
| --- | --- | --- | --- | --- | --- | --- |
| Gleann Dubh,<br>Arran | AR | Scotland | 55.55 | -5.202 | 12 | 11 |
| Blowick Fell | BF | Lake District | 54.558 | -2.931 | 10 | 10 |
| Balnaguard<br>Glen | BG | Scotland | 56.644 | -3.73 | 23 | 30 |
| Bulford Hill | BH | S. England | 51.204 | -1.7 | 12 | 11 |
| Blea Tarn | BT | Lake District | 54.426 | -3.088 | 10 | 8 |
| Clashindarroch | CD | Scotland | 57.338 | -2.967 | 20 | 35 |
| Danebury Hill | DH | S. England | 51.137 | -1.535 | 22 | 27 |
| Fasnakyle | FK | Scotland | 57.336 | -4.849 | 22 | 26 |
| Glen Artney | GA | Scotland | 56.34 | -3.996 | 23 | 24 |
| Invernaver | IN | Scotland | 58.521 | -4.256 | 6 | 6 |
| Lammermuir | LM | Scotland | 55.853 | -2.71 | 23 | 30 |
| Morrone<br>Birkwood | MB | Scotland | 56.998 | -3.426 | 7 | 6 |
| Porton Down | PD | S. England | 51.138 | -1.652 | 22 | 24 |
| Thwaites Fell | TF | Lake District | 54.302 | -3.263 | 22 | 25 |
| Tynron | TY | Scotland | 55.214 | -3.846 | 15 | 15 |
| Whitewell | WW | Scotland | 57.155 | -3.796 | 9 | 9 |
| *ssp. nana* | *Nana* | Scotland | 56.003 | -5.635 | 8 |  |
| *ssp.*<br>*hemisphaerica* | *Hemi* | n/a | 49.961 | -5.213 | 11 | 7 |

Prior to DNA extraction, needles were freeze-dried and finely chopped using a razor blade, and then using a Retsch MM 300 mixer mill at a frequency of 30/s for a total of 2 minutes with metal beads in each Eppendorff tube to grind samples. Razor blades, scissors and lab surfaces were cleaned with 2% ethanol between processing samples. DNA was extracted for each sample using a Qiagen DNeasy Plant Pro kit following the manufacturer’s instructions. SNPs were identified and sequenced for this study by the Plant Genomic Resources Centre (CNRGV, INRAE, France) (https://cnrgv.toulouse.inrae.fr/).

Approximately 90,000 loci were initially evaluated by CNRGV, which were then filtered to return 74 loci for the following analyses. CNRGV sequentially applied the following filters: 1) loci where a genotype was called for all individuals (including technical replicates; 9,539 loci remaining), 2) loci with functional technical replicates (9,374 loci remaining), 3) loci with unique SNPs in the returned sequence (807 loci remaining), 4) loci with at least 3 genotypes (198 loci remaining) and 5) loci where the unique SNP site is not in the first or last 20 bases of the sequence to allow for primer design (175 loci remaining). Of these remaining 175 loci, 80 were selected to develop two multiplex Sequenom SNP chips which provided data on 74 SNP loci in the samples analysed in this study (https://cnrgv.toulouse.inrae.fr/) (Appendix 1).

A new set of microsatellite markers was developed for this study (Appendix 2).

Juniper DNA samples were sent to Microsynth Ecogenics (https://www.ecogenics.ch/home.html) to identify nuclear SSR containing sequences. Of the 285 identified microsatellites 48 were tested using multiplex PCR and gel electrophoresis which returned 6 polymorphic markers which were consistently amplified by PCR. For each of the six loci, PCR was carried out as follows: each forward primer had a 5’ – AGGTTTTCCCAGTCACGACGTT – 3’ M13 sequence attached at the 5’ end for subsequent detection purposes. DNA was amplified in a total volume of 20µl comprising the following reaction mixture: 1.5µl DNA, 1X PCR buffer (Bioron, Germany), 5µM of each primer (0.2mM of each dNTP (VWR International), 0.25µM M13 oligo with a fluorescent dye attached, and 0.25U Taq DNA polymerase (Bioron). PCR products were run on a Licor 4300 DNA sequencer and the allele sizes scored by eye using a size marker as reference.

### Data cleaning

Marker data were cleaned and checked: monomorphic loci, those with more than 10% missing data, and those out of Hardy-Weinburg Equilibrium (HWE) in more than half of all populations were removed using GenAlEx (Peakall & Smouse, 2012). The SSR data were screened for null alleles using Microchecker (Van Oosterhout et al., 2004). Since junipers are dioecous, and the markers were most likely selectively neutral, we expect the loci to be in HWE; either loci out of HWE or null alleles may cause a false heterozygote excess. Finally, genotype data were screened for clonal individuals (population restoration is often done using cuttings) using R (Team, 2021) and the dplyr package (Wickham et al., 2023). One individual was removed when two individuals within a population shared an exact genotype, but when genotypes were shared between individuals in different populations, both were retained for the analyses.

### Data analyses: descriptive statistics and fixation indexes

The following descriptive statistics were calculated for each population using GenAlEx (Peakall & Smouse, 2012): average number of individuals (N), number of alleles (N_a_), observed (H_o_) and expected heterozygosity (H_e_), the inbreeding fixation index (F_is_) and the standard error for the inbreeding fixation index (SE F_is_). Both Wright’s standard fixation index (F_st_) and Wright’s adjusted fixation index (F’_st_) were calculated using GenAlEx (Peakall & Smouse, 2012) both of which are reported here, following the recommendation of Peakall and Smouse (2012). To facilitate comparisons within regions, the arithmetic means of pairwise comparisons between populations for both fixation indices were calculated for all populations within each region.

### Data analyses: Population structure and genetic differentiation

A hierarchical analysis of molecular variance (AMOVA) was performed using GenAlEx (Peakall & Smouse, 2012) with 9999 permutations and within-individual variability suppressed. To facilitate comparisons within and between regions, the arithmetic mean of pairwise fixations indices comparing populations within and between regions were calculated. A Principal Coordinate Analysis (PCoA) in addition to a test for Isolation by Distance (IBD) were both performed using GenAlEx (Peakall & Smouse, 2012). IBD was evaluated by running a Mantel test comparing linear geographic distances in kilometres with pairwise, individual-by individual genetic distances for both all populations and for only Scottish populations due to their larger geographic distribution than the other population centres. We tested for genetic structure using the STRUCTURE software v.2.3.4 (Pritchard et al., 2000). The Markov chain Monte Carlo (MCMC) method used a burn-in length of 10,000 steps, followed by 10,000 steps for estimation. Each simulation was replicated with 20 iterations for each K value between two and 21, from which mean values were calculated.

These data were uploaded to STRUCTURE Harvester (Earl & vonHoldt, 2012) to estimate the most likely K value using delta K method (Evanno et al., 2005) and to CLUMPAK (Kopelman et al., 2015) to summarize the Q matrices for all 20 iterations of each K into a single Q-matrix. The returned Q-matrices for each K value were processed into spatial objects using the raster package for R (Hijmans, 2023) and plotted as pie charts in ArcMap. We report the results for K=3, 4 and 5 for both SNP and SSR datasets.

### Subspecies exclusions

The population of *Hemi* was removed when calculating pairwise F_st_ values and performing the AMOVA, IBD and PCoA analyses for two reasons. Firstly, only 39.2% of SNP and 50% of SSR loci were polymorphic for *Hemi*. Secondly, *Hemi* was highly differentiated from the other populations, with pairwise F_st_ values between *Hemi* and other populations ranging from 0.359 to 0.481 for SNP data and from 0.238 to 0.499 for SSR data. Including *Hemi* in AMOVA, IBD and PCoA therefore obscures finer-scale differences between the other populations. By contrast, *Nana* was included in all calculations. Unlike *Hemi, Nana* had an acceptable percent of polymorphic loci, and excluding *Nana* from pairwise F_st_ values, AMOVA and PCoA resulted in only minor differences in these values and analyses.

## Results

### Data cleaning

Fourteen SNP loci with more than 10% missing data and a further nine loci were removed because they were monomorphic. An additional ten SNP loci were removed for being significantly out of HWE within more than half of all populations, leaving a total of 41 SNP loci that were used in the analyses. Two loci were removed from the SSR dataset due to being null alleles within more than half of all populations, leaving 4 SSR loci for subsequent analyses. Twelve individuals from the SNP dataset were clonal, sharing five genotypes among them. CD, *Nana* and *Hemi* were the only populations which contained clonal duplicates, and none shared a genotype between other populations. In each case, only one individual per genotype was retained for subsequent analyses (Appendix 3). Thirty-four clonal individuals were identified in the SSR dataset sharing 16 genotypes among them.

*Hemi, Nana*, CD, PD and LM had genotypes shared between individuals, and in each case all were removed except one (Appendix 4). Finally, one individual from WW was removed from the SNP dataset due to being an extreme outlier, as it differed in virtually every locus to all other individuals in that population, suggesting the possibility of an extraction or sequencing error. This left 277 and 304 individuals for the analyses of SNP and SSR data, respectively (Table 1).

### Descriptive statistics

In the SNP dataset the percent of polymorphic loci within populations was relatively high, ranging from 68.3% (*Nana*) to 97.6% (CD) with an average of 88.4%. F_is_ values from the SNP dataset were generally low, with an overall mean of 0.025, and a range from −0.221 (*Hemi*) to 0.108 (BF) (Table 2). All populations had 100% polymorphic loci in the SSR dataset except for *Hemi*, which had 50% polymorphic loci. F_is_ values calculated from the SSR dataset were also generally low, with a mean of −0.002 and a range from −0.159 (*Nana*) to 0.171 (BG) (Table 3). Six populations (BG, CD, DH, FK, GA and PD) had positive F_is_ values for both SNP and SSR datasets; eight populations (AR, BF, BH, LM, *Nana*, TF, TY and WW) have one positive value and one negative value between the SNP and SSR datasets; the remaining four populations (BT, *Hemi,* IN and MB) had negative values for both datasets.

**Table 2:** Mean descriptive statistics for SNP data across all 51 loci. Key: N= average number of individuals with valid data per loci, N_a_= number of alleles, H_o_= observed heterozygosity, H_e_=expected heterozygosity, uH_e_=unbiased expected heterozygosity, F_is_= inbreeding fixation index, F_is_ SE=Standard error of fixation index.

| Pop | N | N<sub>a</sub> | H<sub>o</sub> | H<sub>e</sub> | uH<sub>e</sub> | F<sub>is</sub> | F<sub>is</sub> SE | % Polymorphic |
| --- | --- | --- | --- | --- | --- | --- | --- | --- |
| AR | 11.9 | 1.902 | 0.298 | 0.327 | 0.341 | 0.076 | 0.054 | 90.24% |
| BF | 9.9 | 1.927 | 0.244 | 0.292 | 0.307 | 0.108 | 0.054 | 92.68% |
| BG | 22.2 | 1.951 | 0.289 | 0.305 | 0.312 | 0.051 | 0.042 | 95.12% |
| BH | 11.8 | 1.902 | 0.274 | 0.293 | 0.306 | 0.090 | 0.053 | 90.24% |
| BT | 9.9 | 1.854 | 0.279 | 0.277 | 0.292 | -0.001 | 0.054 | 85.37% |
| CD | 19.8 | 1.976 | 0.299 | 0.315 | 0.323 | 0.022 | 0.046 | 97.56% |
| DH | 21.9 | 1.902 | 0.265 | 0.281 | 0.288 | 0.030 | 0.043 | 90.24% |
| FK | 21.8 | 1.951 | 0.267 | 0.288 | 0.295 | 0.074 | 0.045 | 95.12% |
| GA | 22.9 | 1.902 | 0.275 | 0.310 | 0.317 | 0.084 | 0.050 | 90.24% |
| HEM | 10.9 | 1.439 | 0.201 | 0.160 | 0.168 | -0.221 | 0.043 | 43.90% |
| IN | 5.9 | 1.829 | 0.340 | 0.295 | 0.323 | -0.145 | 0.060 | 82.93% |
| LM | 22.8 | 1.902 | 0.291 | 0.297 | 0.304 | 0.020 | 0.042 | 90.24% |
| MB | 6.9 | 1.829 | 0.336 | 0.330 | 0.356 | -0.018 | 0.067 | 82.93% |
| NANA | 8.0 | 1.683 | 0.220 | 0.227 | 0.242 | 0.023 | 0.069 | 68.29% |
| PD | 21.9 | 1.878 | 0.259 | 0.295 | 0.302 | 0.078 | 0.038 | 87.80% |
| TF | 21.5 | 1.902 | 0.286 | 0.268 | 0.275 | -0.046 | 0.040 | 90.24% |
| TY | 14.7 | 1.927 | 0.252 | 0.267 | 0.276 | 0.033 | 0.050 | 92.68% |
| WW | 8.9 | 1.805 | 0.263 | 0.277 | 0.293 | 0.044 | 0.066 | 80.49% |
| Mean | 15.2 | 1.859 | 0.274 | 0.284 | 0.296 | 0.025 | n/a | 85.91% |
| SE | 0.23 | 0.013 | 0.007 | 0.006 | 0.007 | 0.012 | n/a | 2.94% |

**Table 3:** Mean descriptive statistics for SSR data across all 5 loci. Key: N= number of individuals, N_a_= number of alleles, H_o_= observed heterozygosity, H_e_=expected heterozygosity, uH_e_= unbiased expected heterozygosity, F_is_=Fixation index, F_is_ SE= standard error for fixation index.

| Pop | N | N<sub>a</sub> | H<sub>o</sub> | H<sub>e</sub> | uH<sub>e</sub> | F<sub>is</sub> | F<sub>is</sub> SE |
| --- | --- | --- | --- | --- | --- | --- | --- |
| AR | 11 | 5.000 | 0.614 | 0.574 | 0.602 | -0.074 | 0.009 |
| BF | 10 | 5.000 | 0.700 | 0.646 | 0.680 | -0.099 | 0.166 |
| BG | 30 | 6.750 | 0.558 | 0.655 | 0.666 | 0.171 | 0.090 |
| BH | 11 | 4.750 | 0.568 | 0.548 | 0.574 | -0.068 | 0.099 |
| BT | 8 | 4.250 | 0.563 | 0.557 | 0.594 | -0.025 | 0.092 |
| CD | 35 | 7.500 | 0.586 | 0.608 | 0.617 | 0.051 | 0.094 |
| DH | 27 | 6.000 | 0.574 | 0.663 | 0.675 | 0.158 | 0.069 |
| FK | 26 | 6.500 | 0.558 | 0.610 | 0.622 | 0.077 | 0.068 |
| GA | 24 | 7.250 | 0.635 | 0.678 | 0.692 | 0.060 | 0.046 |
| HEM | 7 | 2.000 | 0.321 | 0.281 | 0.302 | -0.117 | 0.184 |
| IN | 6 | 3.250 | 0.500 | 0.510 | 0.557 | -0.003 | 0.119 |
| LM | 30 | 5.000 | 0.608 | 0.582 | 0.592 | -0.065 | 0.046 |
| MB | 6 | 4.500 | 0.750 | 0.688 | 0.750 | -0.105 | 0.218 |
| NANA | 6 | 3.750 | 0.625 | 0.507 | 0.553 | -0.159 | 0.137 |
| PD | 24 | 5.750 | 0.594 | 0.658 | 0.672 | 0.098 | 0.049 |
| TF | 25 | 5.250 | 0.560 | 0.582 | 0.594 | 0.007 | 0.081 |
| TY | 15 | 4.000 | 0.583 | 0.588 | 0.609 | -0.018 | 0.080 |
| WW | 9 | 4.750 | 0.611 | 0.603 | 0.639 | -0.032 | 0.096 |
| Mean | 17.2 | 5.069 | 0.584 | 0.585 | 0.610 | -0.002 | n/a |
| SE | 1.1 | 0.260 | 0.029 | 0.026 | 0.027 | 0.024 | n/a |

### Fixation indices

Fixation indices were slight but significant among most S. English and Scottish populations, but nonsignificant among the Lake District populations in both datasets, indicating minor but detectable genetic differences among populations within S. England and Scotland and a lack of genetic differentiation among the Lake District populations. The overall average fixation index (F_st_) for SNP data within S. English, Lake District and Scottish populations were 0.029, 0.017 and 0.050, with ranges from 0.028 (BH-PD and DH-PD) to 0.031 (DH-BH), 0.011 (BT-TF) to 0.022 (BT-BF) and 0 (MB-GA) to 0.134 (*Nana*-WW), respectively. The overall average of the adjusted fixation index (F’st) within S. English, Lake District and Scottish populations were 0.041, 0.025 and 0.072 with ranges from 0.040 (BH- PD, DH-PD) to 0.044 (BH-DH), 0.016 (BF-TF) to 0.031 (BF-BT) and −0.009 (GA-MB) to 0.185 (*Nana*-WW), respectively (Tables 4-6).

**Table 4:** Pairwise F_st_ values (below diagonal) with stars signifying significance levels and adjusted F’_st_ values (above diagonal) for S. English populations from SNP dataset. * signifies p value between 0.05-0.005.

|  |  |  |  |
| --- | --- | --- | --- |
| BH |  | 0.044 | 0.040 |
| DH | 0.0306* |  | 0.040 |
| PD | 0.0277* | 0.028* |  |
|  | BH | DH | PD |

**Table 5:** Pairwise F_st_ values (below diagonal) levels and adjusted F’_st_ values (above diagonal) for Lake District populations from SNP dataset. F_st_ values were nonsignificant.

|  |  |  |  |
| --- | --- | --- | --- |
| BF |  | 0.031 | 0.027 |
| BT | 0.022 |  | 0.016 |
| TF | 0.019 | 0.011 |  |
|  | BF | BT | TF |

**Table 6:**
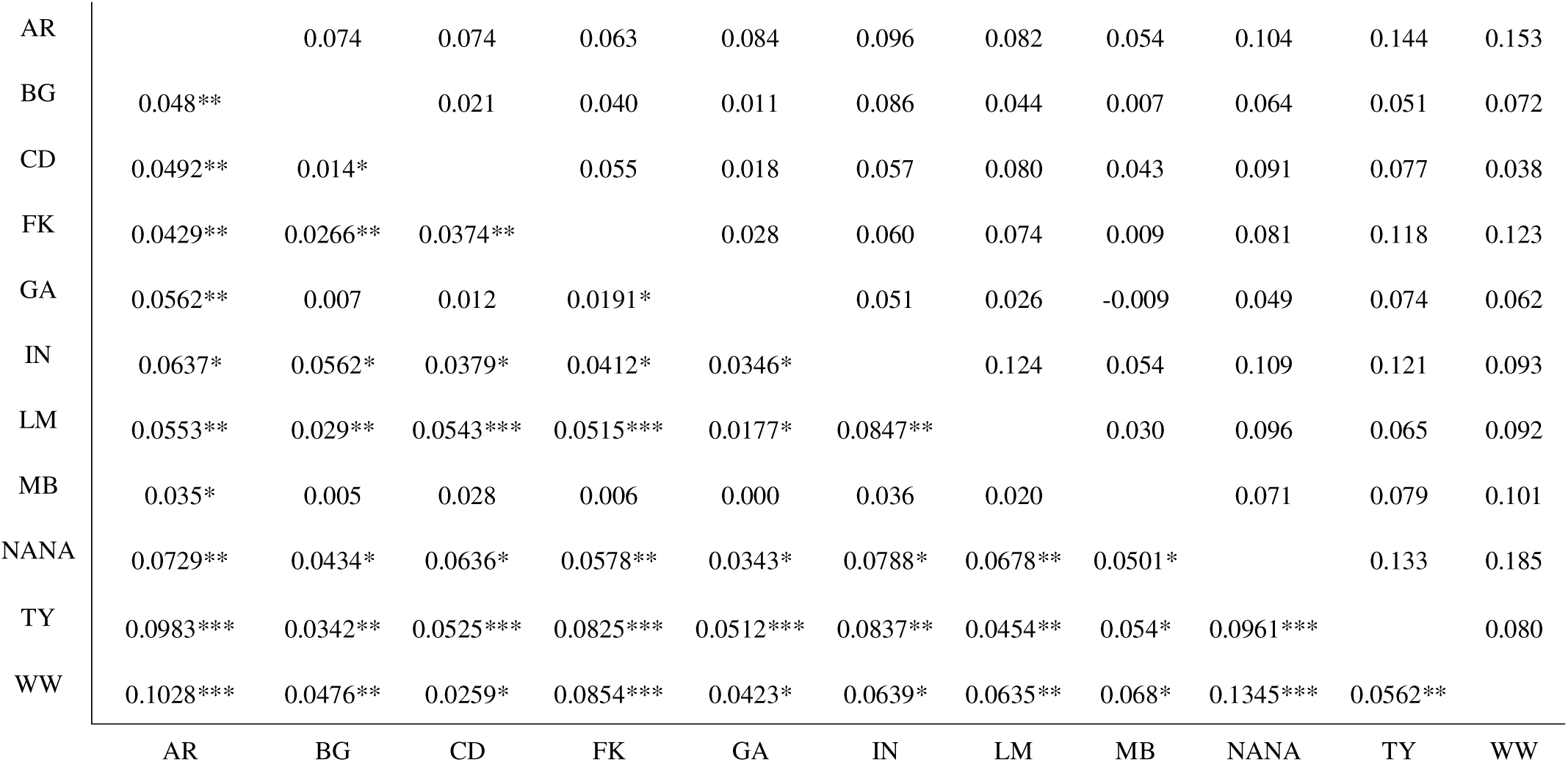
Pairwise F_st_ values (below diagonal) with stars signifying significance levels and adjusted F’_st_ values (above diagonal) for Scottish populations from SNP dataset. * signifies p value between 0.05-0.005; ** signifies p value between 0.0049-0.0005, *** signifies p value between 0.00049-0.000005

|  |  |  |  |  |  |  |  |  |  |  |  |
| --- | --- | --- | --- | --- | --- | --- | --- | --- | --- | --- | --- |
| AR |  | 0.074 | 0.074 | 0.063 | 0.084 | 0.096 | 0.082 | 0.054 | 0.104 | 0.144 | 0.153 |
| BG | 0.048** |  | 0.021 | 0.040 | 0.011 | 0.086 | 0.044 | 0.007 | 0.064 | 0.051 | 0.072 |
| CD | 0.0492** | 0.014* |  | 0.055 | 0.018 | 0.057 | 0.080 | 0.043 | 0.091 | 0.077 | 0.038 |
| FK | 0.0429** | 0.0266** | 0.0374** |  | 0.028 | 0.060 | 0.074 | 0.009 | 0.081 | 0.118 | 0.123 |
| GA | 0.0562** | 0.007 | 0.012 | 0.0191* |  | 0.051 | 0.026 | -0.009 | 0.049 | 0.074 | 0.062 |
| IN | 0.0637* | 0.0562* | 0.0379* | 0.0412* | 0.0346* |  | 0.124 | 0.054 | 0.109 | 0.121 | 0.093 |
| LM | 0.0553** | 0.029** | 0.0543*** | 0.0515*** | 0.0177* | 0.0847** |  | 0.030 | 0.096 | 0.065 | 0.092 |
| MB | 0.035* | 0.005 | 0.028 | 0.006 | 0.000 | 0.036 | 0.020 |  | 0.071 | 0.079 | 0.101 |
| NANA | 0.0729** | 0.0434* | 0.0636* | 0.0578** | 0.0343* | 0.0788* | 0.0678** | 0.0501* |  | 0.133 | 0.185 |
| TY | 0.0983*** | 0.0342** | 0.0525*** | 0.0825*** | 0.0512*** | 0.0837** | 0.0454** | 0.054* | 0.0961*** |  | 0.080 |
| WW | 0.1028*** | 0.0476** | 0.0259* | 0.0854*** | 0.0423* | 0.0639* | 0.0635** | 0.068* | 0.1345*** | 0.0562** |  |
|  | AR | BG | CD | FK | GA | IN | LM | MB | NANA | TY | WW |

For SSR data, the overall average fixation index (F_st_) within S. English, Lake District and Scottish populations were 0.025, 0.013 and 0.063 with ranges from 0.006 (BH-PD) to 0.053 (BH-DH), 0 (BF-TF) to 0.21 (BT-BF) and 0.001 (MB-WW) to 0.137 (LM-IN), respectively. The overall average of the adjusted fixation index (F’_st_) for SSR data within S. English, Lake District and Scottish populations were 0.069, 0.035 and 0.165 with ranges from 0.018 (BH-PD) to 0.144 (BH-DH), −0.001 (BF-TF) to 0.059 (BF-BT) and 0.002 (MB-WW) to 0.327 (IN-LM), respectively (Tables 7-9).

**Table 7:**
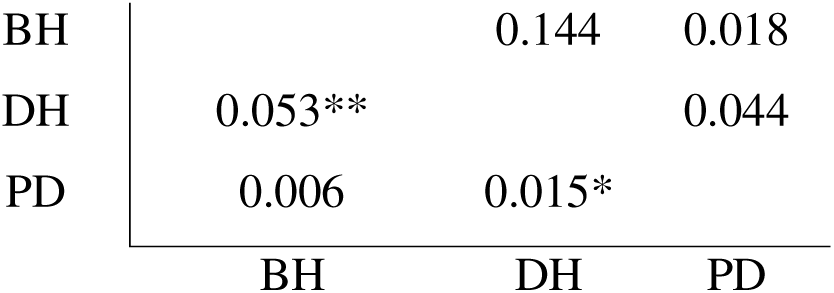
Pairwise F_st_ values (below diagonal) with stars signifying significance levels and adjusted F’_st_ values (above diagonal) for S. English populations from SSR dataset. * signifies p value between 0.05-0.005, ** signifies p value between 0.0049-0.0005.

**Table 8:**
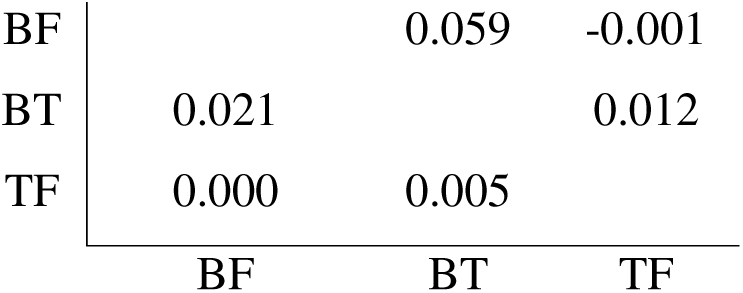
Pairwise F_st_ values (below diagonal) and adjusted F’_st_ values (above diagonal) for Lake District populations from SSR dataset. F_st_ values were nonsignificant.

|  |  |  |  |
| --- | --- | --- | --- |
| BF |  | 0.059 | -0.001 |
| BT | 0.021 |  | 0.012 |
| TF | 0.000 | 0.005 |  |
|  | BF | BT | TF |

**Table 9:** Pairwise F_st_ values (below diagonal) with stars signifying significance levels and adjusted F’_st_ values (above diagonal) for Scottish populations from SSR dataset. * signifies p value between 0.05-0.005; ** signifies p value between 0.0049-0.0005, *** signifies p value between 0.00049-0.000005

|  |  |  |  |  |  |  |  |  |  |  |  |
| --- | --- | --- | --- | --- | --- | --- | --- | --- | --- | --- | --- |
| AR |  | 0.198 | 0.110 | 0.183 | 0.178 | 0.257 | 0.277 | 0.177 | 0.259 | 0.245 | 0.284 |
| BG | 0.071*** |  | 0.049 | 0.086 | 0.034 | 0.290 | 0.098 | 0.141 | 0.072 | 0.190 | 0.087 |
| CD | 0.043** | 0.018* |  | 0.094 | 0.025 | 0.196 | 0.108 | 0.086 | 0.077 | 0.117 | 0.122 |
| FK | 0.071*** | 0.031*** | 0.036*** |  | 0.070 | 0.233 | 0.218 | 0.166 | 0.081 | 0.270 | 0.195 |
| GA | 0.062*** | 0.011 | 0.009 | 0.024* |  | 0.211 | 0.137 | 0.037 | 0.097 | 0.162 | 0.107 |
| IN | 0.107** | 0.108*** | 0.079** | 0.093** | 0.076** |  | 0.327 | 0.243 | 0.241 | 0.095 | 0.267 |
| LM | 0.112*** | 0.037** | 0.043*** | 0.086*** | 0.049*** | 0.137*** |  | 0.233 | 0.221 | 0.121 | 0.141 |
| MB | 0.059* | 0.043* | 0.029 | 0.055* | 0.010 | 0.084* | 0.082** |  | 0.281 | 0.167 | 0.002 |
| NANA | 0.109*** | 0.027 | 0.031 | 0.033 | 0.035* | 0.107* | 0.093** | 0.098* |  | 0.237 | 0.277 |
| TY | 0.097*** | 0.068*** | 0.045*** | 0.104*** | 0.056*** | 0.039 | 0.048** | 0.056* | 0.098** |  | 0.178 |
| WW | 0.108*** | 0.03* | 0.046* | 0.073*** | 0.035* | 0.106** | 0.055** | 0.001 | 0.111** | 0.067** |  |
|  | AR | BG | CD | FK | GA | IN | LM | MB | NANA | TY | WW |

### Fixation indices within and between regions

Regions showed markedly different fixation index values, with comparisons between populations in Scotland and S. England having the highest values in the SNP dataset (F_st_=0.095 and F’_st_=0.137) and those between Scotland and the Lake District (F_st_=0.136 and F’_st_=0.231) having the highest values in the SSR dataset. In both datasets, comparisons within the Lake District and S. England were smaller than those within Scotland. The average of all pairwise F_st_ and F’_st_ values comparing populations within and between regions for SNP data ranged from 0.017 to 0.095 (F_st_, Lake District- Lake District and Scotland- S. England, respectively) and 0.025-0.137 (F’_st_, Lake District- Lake District and Scotland-S. England, respectively) (Table 10). For SSR data, these comparisons varied from 0.027 to 0.136 (F_st_, Lake District- Lake District and Scotland- Lake District, respectively) and 0.023-0.231 (F’_st_, Lake District- Lake District and Scotland-Lake District, respectively) (Table 11).

**Table 10:** Average of pairwise F_st_ (top) and adjusted F’_st_ (bottom) values among all populations within and between the three regions for SNP data.

| Fst | Scotland | Lake District | S. England |
| --- | --- | --- | --- |
| Scotland | 0.050 |  |  |
| Lake District | 0.064 | 0.017 |  |
| S. England | 0.095 | 0.081 | 0.029 |
| F'st | Scotland | Lake District | S. England |
| Scotland | 0.072 |  |  |
| Lake District | 0.093 | 0.025 |  |
| S. England | 0.137 | 0.117 | 0.041 |

**Table 11:** Average of pairwise F_st_ (top) and adjusted F’_st_ (bottom) values among all populations within and between the three regions for SSR data.

| Fst | Scotland | Lake District | S. England |
| --- | --- | --- | --- |
| Scotland | 0.067 |  |  |
| Lake District | 0.136 | 0.045 |  |
| S. England | 0.079 | 0.044 | 0.027 |
| F'st | Scotland | Lake District | S. England |
| Scotland | 0.165 |  |  |
| Lake District | 0.231 | 0.023 |  |
| S. England | 0.163 | 0.210 | 0.069 |

### Population structure: AMOVA

The hierarchical AMOVA’s for both SNP and SSR data ascribed a large majority of the observed genetic variance to that within a population for both SNP and SSR data (92% for both SNP and SSR datasets). The variance between populations was significant for both SNP (F_st_=0.079; p<0.001) and SSR data (F_st_=0.076; p<0.001). The variance between regions was also significant for both SNP (F_rt_=0.043; p<0.001) and SSR data (F_rt_=0.037; p<0.001).

### Population structure: IBD

There were highly significant positive correlations between geographic and genetic distances over all populations in both SNP (R^2^=0.018, p=0.0001) and SSR (R^2^=0.045, p=0.005) datasets. However, within just Scottish populations neither SNP (R^2^=0.001, p=0.190) nor SSR (R^2^=0.0001, p=0.366) datasets detected significant IBD.

### Population structure: PCoA

The first two principal coordinates cumulatively accounted for 46.75 and 50.11 percent of the observed variation for SNP and SSR datasets, respectively (Tables 12 and 13). In both datasets, populations within each region clustered together. The groupings are especially pronounced in the SNP dataset (Figure 3). In Figure 4, the two principal coordinate axes are switched because they describe a similar percentage of the variation (Table 13) and to provide better visual alignment with Figure 3.

**Figure 3:**
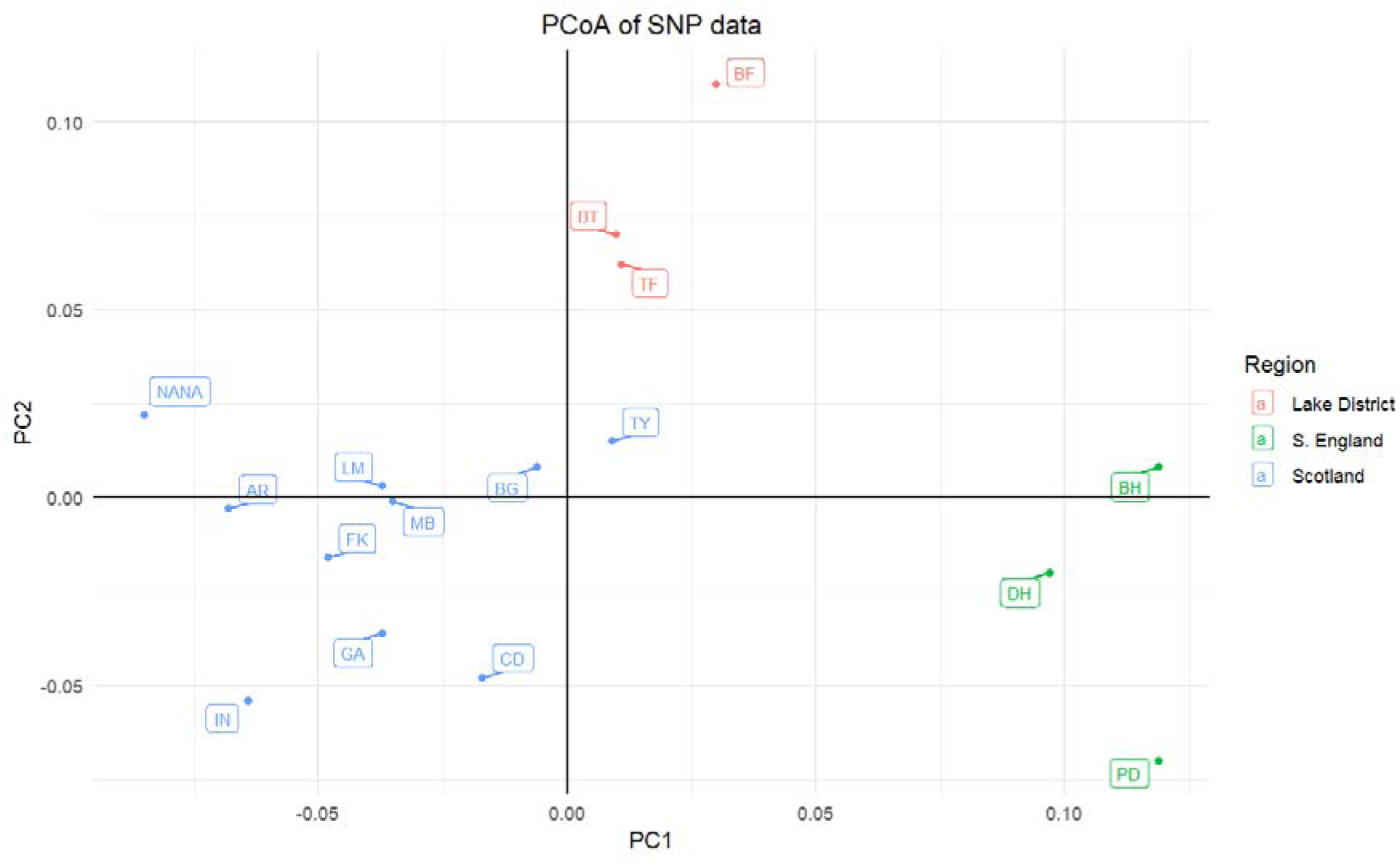
Principal coordinate analysis (PCoA) of SNP dataset. Populations are labelled with the abbreviations in table 1 and regions are colour coded by region.

**Figure 4:**
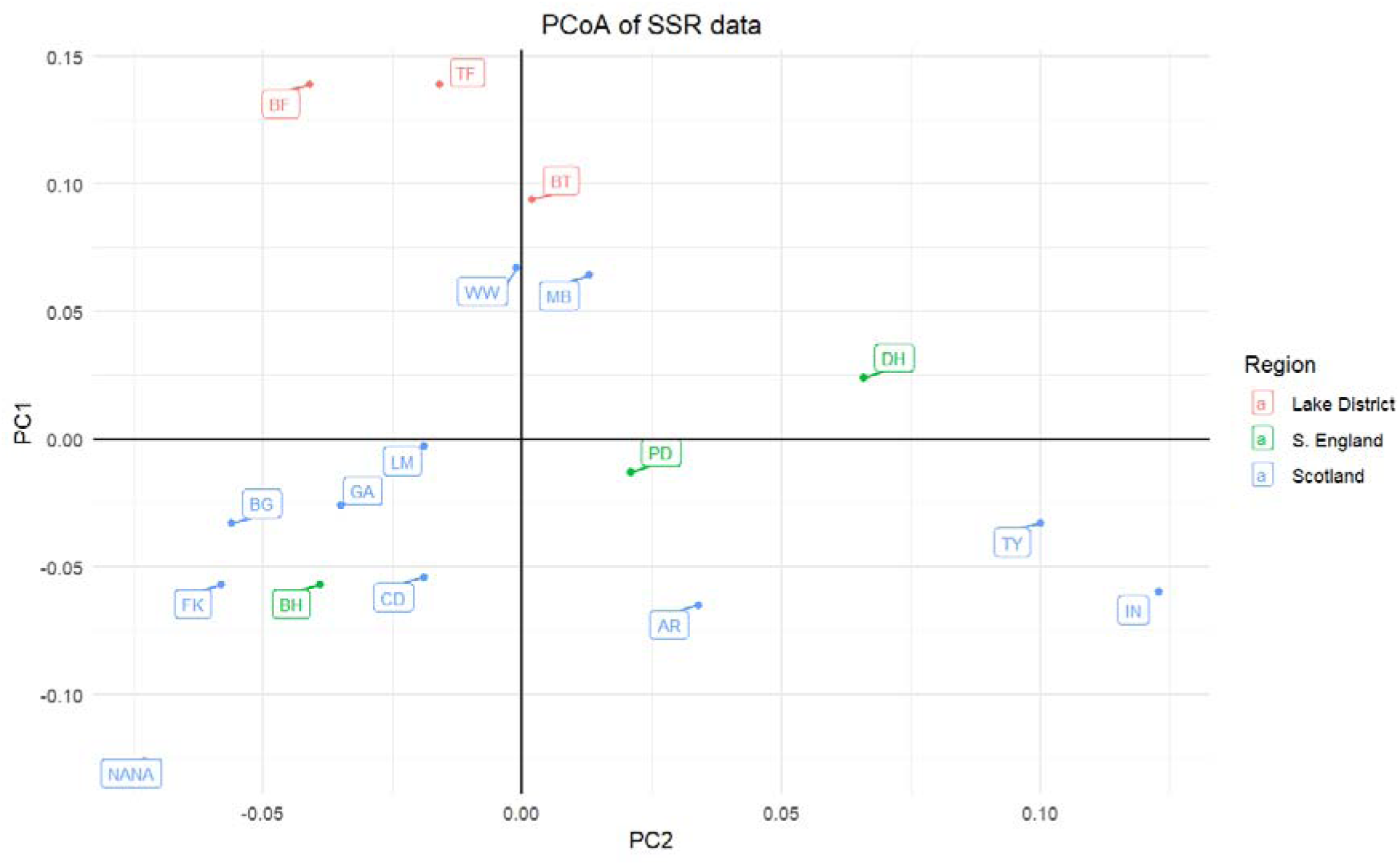
Principal coordinate analysis (PCoA) of SSR dataset. Populations are labelled with the abbreviations in table 1 and regions are colour coded by region.

**Table 12:** Percent of observed genetic variation explained by each of first three axes, and cumulative percentage explained for SNP data.

| Axis | 1 | 2 | 3 |
| --- | --- | --- | --- |
| Percent variation explained | 28.48 | 18.27 | 14.19 |
| Cumulative percent variation explained | 28.48 | 46.75 | 60.94 |

**Table 13:**
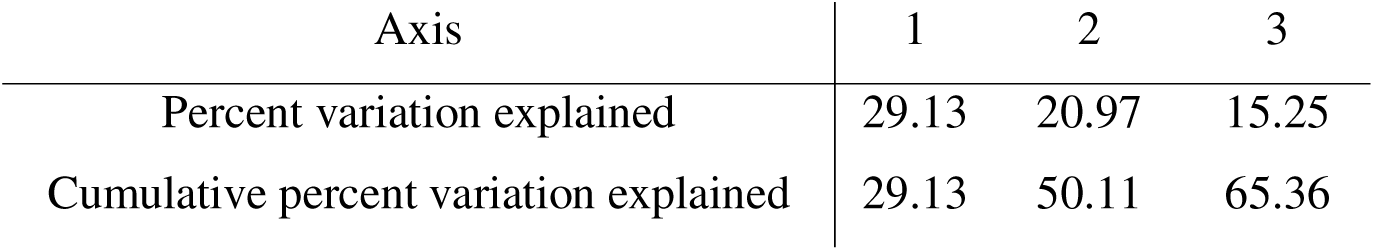
Percent of observed genetic variation explained by each of first three axes, and cumulative percentage explained for SSR data.

| Axis | 1 | 2 | 3 |
| --- | --- | --- | --- |
| Percent variation explained | 29.13 | 20.97 | 15.25 |
| Cumulative percent variation explained | 29.13 | 50.11 | 65.36 |

### Population structure: STRUCTURE and CLUMPAK

In our tests for population structure, we found that the optimal Delta K for the SNP dataset was K=4, and for the SSR dataset the optimal Delta K is K=3, with K=5 as the second highest value (Figure 5). In both datasets, the S. England populations group strongly together and the population of *Hemi* is ascribed almost entirely to a single genetic group. Lake District populations are grouped together and mostly exclude populations from the other regions at K=5 (SNP) and K=4 (SSR). Scottish populations showed some sign of sub-structuring into two groups, particularly in the SSR dataset, with those populations in the Highlands (IN, FK, CD, WW, MB, GA, AR) and those in the Borders (TY, LM) grouping together, although both genetic groups were present throughout. This pattern is notable at SNP K=5 and at SSR K=3-5.

**Figure 5:**
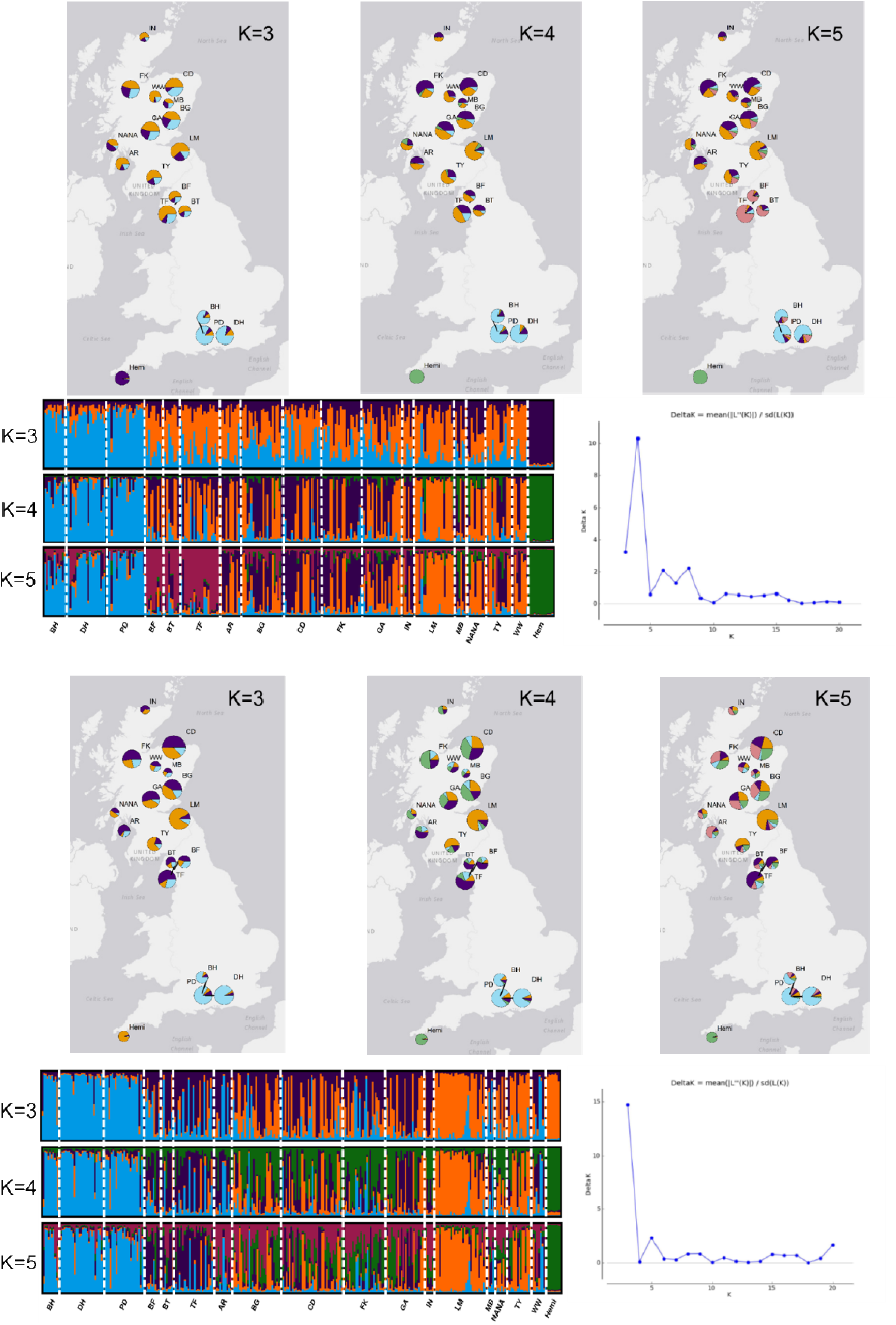
Results from STRUCTURE runs based on SNP (top) SSR (bottom) datasets for K=3, K=4 and K=5. Pie charts display locations of sampled populations and proportional membership of each population to each genetic group, with the size of each chart being proportional to the number of sampled individuals from that population. Bar graphs display proportional membership of each individual to each genetic group, where each bar is an individual. Colours represent genetic groups; white dashed lines separate populations, which are specified along the x-axis.

## Discussion

Our analyses of two types of neutral genetic markers in UK juniper populations support the existence of at least three distinct genetic groups within *J. communis*: S. English populations, Lake District populations and Scottish populations. In addition, F_st_ values and STRUCTURE analyses suggested a potential fourth genetic group in the Scottish Borders. The three genetic groups have different patterns of genetic diversity and likely experience different amounts of gene flow among their respective populations. Our analyses also indicated a clear genetic distinctiveness of *Hemi* from other populations of *J. communis* and *Nana,* which supports previous studies that describe *Hemi* as a separate subspecies (Thomas et al., 2007).

Our findings are consistent with previous studies of juniper population genetics on the British Isles in that they have all found evidence of significant population differentiation and limited gene flow between remnant populations (Merwe et al., 2000; Provan et al., 2008; Reynolds, 2022). Advancing on this knowledge, we have brought together population-level samples from across Great Britain. Our results indicate that habitat fragmentation is having a notable negative impact, causing barriers to gene flow among UK juniper stands and inbreeding within some of the sampled populations.

García et al. (2018) evaluated the performance of SNP versus SSR markers in Mediterranean populations of *J. phoenicea ssp. turbinata*. Their work demonstrated that SSR markers generally have both larger values and variation in estimating diversity indices, making SNP markers better suited for estimating genome-wide genetic diversity and inferring historic demographic processes. However, SSR markers outperformed SNP markers in determining parentage (García et al., 2018). In our data, SSR markers returned higher diversity estimates, but they were nonetheless largely consistent with the results from the SNP dataset.

The results of the AMOVA and pairwise, average F_st_ and F’_st_ values by region (Tables 10 and 11) suggest that both regions and almost all populations are significantly distinct from each other, with the exception of Lake District populations, which were not significantly different from one another. These findings suggest restricted gene flow between populations and is consistent with the expectations for a fragmented species (Provan et al., 2008; RobledoLArnuncio et al., 2005). Although the slight to moderate F_st_ and F’_st_ values might suggest that some gene flow still occurs between the three regions, it is possible that this is a reflection of the longevity of juniper trees, meaning that the sampled individuals may be representative of a historically larger metapopulation that has since become fragmented.

The three UK regions had different patterns of genetic diversity between their respective populations, indicated by differences in their average, pairwise F_st_ and F’_st_ values and population-wise F_is_ values (Tables 2-9). Lake District populations, which have nonsignificant F_st_ values and generally slightly negative F_is_ values (Tables 2, 3, 5 and 8) have been or are likely capable of gene flow between populations that balances genetic drift. Southern English populations were very similar to one another (Tables 2, 3, 4 and 7), but two of the three were nonetheless significantly differentiated by genetic markers and all had positive F_is_ values (with the exception of BH in the SSR dataset). In contrast to the Lake District populations, our data suggest that S. English populations are likely less capable of inter-population gene flow, despite the relatively small geographic area, and furthermore they are likely already experiencing inbreeding and loss of heterozygosity because of this. Scottish populations are generally more differentiated from one another than populations within either of the other regions are to each other (Tables 2, 3, 6 and 9). To some extent this is to be expected, given the larger geographic distances separating the sampled populations and the greater number of sampled populations from Scotland compared to the other regions. Nonetheless, the higher diversity and differentiation of Scottish populations, the significant AMOVA results and the patterns of F_st_, F’_st_ and F_is_ are indicative of a fragmented landscape with substantial barriers to gene flow between populations within Scotland (Oostermeijer & De Knegt, 2004; Provan et al., 2008; Reim et al., 2016). F_is_ values for Scottish populations are highly variable, ranging from 0.171 (BG; SSR dataset) to −0.145 (IN; SNP dataset). F_is_ values do not appear to follow a geographic pattern within Scotland, and the degree of inbreeding in Scottish populations is likely to be situationally dependent.

Three populations had particularly negative F_is_ values in either SNP and/or SSR datasets, indicating heterozygote excess: *Hemi* (negative F_is_ in both both SNP and SSR datasets)*, Nana* (negative F_is_ in SSR dataset) and IN (negative F_is_ in both SNP and SSR datasets). In all three cases, it is probable that this excess is caused by sampling effects (e.g.: artificial propagation of cuttings causing some genotypes to be over-represented in a population), a population expansion following a bottleneck, and/or immigrating individuals from other populations. However, these hypotheses are beyond the scope of the present paper and will be investigated in a future project.

The slight, but significant IBD when including all regions but lack of significant IBD within Scotland is likely due to the patchy distribution of juniper within the UK: S. English populations are very isolated from the other population centres compared to those in Scotland and the UK is a heavily fragmented landscape. It follows that gene flow will be locally biased in this case. The fact that Scottish populations do not have significant IBD indicates that gene flow among Scottish populations is likely not biased in any particular direction. In other words, two Scottish populations that are close together are not statistically likely to be more similar to each other than they are to more distant populations (Meirmans, 2012). This may be reflective of the diversity of the meta-population before fragmentation, especially given the longevity of juniper trees, but it does hint that juniper may be capable of gene flow over considerable distances. Previous literature describing juniper pollen and seed dispersal generally find local deposition (within tens of metres) is most common (Adams & Thornburg, 2010; García, 2001; Surso, 2018), however Hall (1990) reported the long range dispersal of juniper pollen as far as 180km, crossing East-West over the Sangre de Cristo mountains in New Mexico. The lack of IBD between Scottish populations hints that long-distance gene dispersal may occur, but not commonly enough to prevent the genetic differentiation of populations.

Although direct comparisons between population genetic studies using different marker types and sampling regimes is not possible, there is a shared pattern in the population genetics of European *J. communis.* Previous studies on English and Welsh (Merwe et al., 2000), Welsh (Reynolds, 2022) and Irish (Provan et al., 2008) populations have consistently found significant geographic structuring and evidence for the lack of gene flow between remnant populations. By contrast, studies on mainland Europe have generally found low levels of genetic differentiation without a geographic pattern to the observed genetic diversity in the main range of juniper in western Europe (Hantemirova et al., 2012; Hantemirova et al., 2017; Knyazeva & Hantemirova, 2020; Oostermeijer & De Knegt, 2004; Reim et al., 2016; Vanden-Broeck et al., 2011). Two notable exceptions to this pattern are the significant differentiation of populations in Russia’s far east and Caucasus from Western European ones (Hantemirova et al., 2012; Hantemirova et al., 2017; Knyazeva & Hantemirova, 2020), and the high degree of differentiation, but lack of geographic signals, that Michalczyk et al. (2010) reported between populations from across central Europe. These studies may be highlighting the geographic distances that juniper gene flow can span: the latter studies, which found geographic patterns in population differentiation, did so over larger areas, whereas those that don’t find patterns in population differentiation are generally across smaller areas.

Juniper is a pioneer species that can thrive in tundra ecosystems (Hantemirova et al., 2017; Michalczyk et al., 2010; Unterholzner et al., 2020) and is therefore often studied to discern biogeography of hardy plant species after the Last Glacial Maximum (LGM) (Hantemirova et al., 2017; Michalczyk et al., 2010). The observed pattern of Mainland European juniper populations having very little genetic differentiation and a lack of geographic signals is hypothesized to be due to repeated fragmentation and expansion events from glacial refugia across Europe, allowing for gene flow between isolation events or even during glacial events to homogenize mainland populations (Hantemirova et al., 2012; Hantemirova et al., 2017; Knyazeva & Hantemirova, 2020; Michalczyk et al., 2008; Michalczyk et al., 2010). By contrast, the high differentiation that studies from the British Isles report are most often interpreted only as the evidence that barriers to gene flow exist between populations (Merwe et al., 2000; Provan et al., 2008; Reynolds, 2022), and only Merwe et al. (2000) has posited a hypothesis on how England was recolonized by junipers after the Last Glacial Maximum, which the findings of Reynolds (2022) did not support.

Although shedding light on the question of juniper’s biogeography in the UK is beyond the scope of the present study, am ongoing project will evaluate it by including more populations from the UK than the present study in addition to populations from across Eurasia. That project will attempt to answer whether the British Isles were colonized in either a single or multiple events and may be able to identify source populations for those colonisations.

Understanding the current genetic diversity of a species is the first step in dynamic conservation. This study has described the current genetic status of juniper populations in Britain. Our findings suggest that *J. communis* populations in Britain are differentiable into at least three distinct genetic groups, possibly four. Southern English populations have been especially impacted by the current habitat fragmentation, being both differentiated from each other in addition to being slightly inbred. By contrast, the populations in the Lake District seem capable of some degree of gene flow, and those within Scotland are generally more differentiated than those in either of the other regions, indicating the likelihood of both sub- population structuring and limited gene flow between populations.

### Conservation recommendations

Our paper demonstrates the effects that over a century of population decline and habitat fragmentation have had on remnant juniper stands. Although there is evidence of some degree of natural resistance (Green et al., 2020), *Phytophthora austrocedri* remains a primary threat to the longevity of junipers in Britain (Green et al., 2015). This threat is compounded by the genetic isolation and inbreeding of juniper stands, which may make populations less able to adapt and develop resistance to the invasive pathogen. Generally, the best way to conserve junipers is to ensure that they have healthy population sizes and can regenerate naturally to express whatever adaptive potential is present and to track changing conditions, including novel pathogens. Protecting natural stands from overgrazing and creating suitable micro-habitats for their seedlings to establish are likely the most appropriate ways to do so (De Frenne et al., 2020; Verheyen et al., 2005). Planting new material within 2km of existing populations is strongly discouraged because of the associated biosecurity risks (Donald et al., 2021).

At least half of the populations sampled in this study displayed some signs of inbreeding, and almost all of them were genetically differentiated from each other, suggesting that gene flow among populations is limited. Although planting operations near existing populations should be avoided, the establishment of “satellite” populations – small, planted populations interspersed among remnant fragments – may help to overcome these barriers and reconnect populations, thereby facilitating the adaptive potential and resilience of those populations. Such populations should be composed of individuals raised, ideally on-site, under strict biosecurity protocols from locally sourced seeds to minimize the potential risks of inbreeding and environmental mismatch, which Baker *et al*. (2024; in preparation), demonstrates may be a possibility. Future research should focus on the dynamics of pollen and seed dispersal, specifically how best to use satellite plantings to help to reconnect population fragments. Incorporating genetics and conservation data can provide insights on the health of juniper populations, and a focus on conserving not only the trees but also the genes can benefit the adaptability and resilience of populations.

## Data accessibility statement

The genotypic data used in this paper are currently being archived at the Environmental Information Data Centre, and will be published open-access with an associated DOI.

## Author contributions

**James Baker:** Data curation (lead), formal analysis (lead), visualization (lead), writing- original draft (lead), writing- review and editing (equal). **Joan Cottrell:** Conceptualization (equal), project administration (equal), writing- review and editing (equal), supervision (equal). **Richard Ennos:** Conceptualization (equal), project administration (equal), writing- review and editing (equal), supervision (equal). **Annika Perry:** Project administration (equal). **Stuart A’Hara:** Sample sequencing and lab work (lead). **Sara Green:** Sample collection (lead), project administration (equal). **Stephen Cavers:** Conceptualization (lead), project administration (lead), writing- review and editing (equal), supervision (lead).

### Acknowledgements

The authors are very grateful to the funders of this project, The Scottish Forestry Trust, Forest Research, The Botanist Foundation, The Woodland Trust and the UKCEH.

## Works Cited

1. Adams, R. P., & Thornburg, D. (2010). Seed dispersal in Juniperus: A review. Phytologia, 92(2), 424–434.

2. Aguilar, R., Quesada, M., Ashworth, L., Herrerias-Diego, Y., & Lobo, J. (2008). Genetic consequences of habitat fragmentation in plant populations: susceptible signals in plant traits and methodological approaches. Molecular Ecology, 17(24), 5177–5188. 10.1111/j.1365-294X.2008.03971.x

3. Al-Snafi, A. E. (2018). Medical importance of Juniperus communis - a review. Indo American Journal of Pharmaceutical Sciences, 05(05). 10.5281/zenodo.1210529

4. Carrer, M., Pellizzari, E., Prendin, A. L., Pividori, M., & Brunetti, M. (2019). Winter precipitation - not summer temperature - is still the main driver for Alpine shrub growth. Sci Total Environ, 682, 171–179. 10.1016/j.scitotenv.2019.05.152

5. Cavender-Bares, J., & Ramírez-Valiente, J. A. (2017). Physiological Evidence from Common Garden Experiments for Local Adaptation and Adaptive Plasticity to Climate in American Live Oaks (Quercus Section Virentes): Implications for Conservation Under Global Change. In E. Gil-Pelegrín, J. J. Peguero-Pina, & D. Sancho-Knapik (Eds.), Oaks physiological ecology. exploring the functional diversity of genus quercus L. (Vol. 7, pp. 107–135). Springer International Publishing. 10.1007/978-3-319-69099-5_4

6. Cavers, S., & Cottrell, J. E. (2015). The basis of resilience in forest tree species and its use in adaptive forest management in Britain. Forestry, 88(1), 13–26. 10.1093/forestry/cpu027

7. Chybicki, I. J., Robledo-Arnuncio, J. J., Bodziarczyk, J., Widlak, M., Meyza, K., Oleksa, A., & Ulaszewski, B. (2024). Disrupted connectivity within a metapopulation of a wind-pollinated declining conifer, Taxus baccata L. Forest Ecosystems, 100240. 10.1016/j.fecs.2024.100240

8. De Frenne, P., Gruwez, R., Hommel, P. W. F. M., De Schrijver, A., Huiskes, R. P. J., De Waal, R. W., Vangansbeke, P., & Verheyen, K. (2020). Effects of heathland management on seedling recruitment of common juniper (Juniperus communis). Plant ecology and evolution, 153(2), 188–198. 10.5091/plecevo.2020.1656

9. De Kort, H., Vandepitte, K., & Honnay, O. (2013). A meta-analysis of the effects of plant traits and geographical scale on the magnitude of adaptive differentiation as measured by the difference between QST and FST. Evolutionary Ecology, 27(6), 1081–1097. 10.1007/s10682-012-9624-9

10. de Villemereuil, P., Gaggiotti, O. E., Mouterde, M., & Till-Bottraud, I. (2016). Common garden experiments in the genomic era: new perspectives and opportunities. Heredity, 116(3), 249–254. 10.1038/hdy.2015.93

11. Dobeš, C., Konrad, H., & Geburek, T. (2017). Potential Population Genetic Consequences of Habitat Fragmentation in Central European Forest Trees and Associated Understorey Species—An Introductory Survey. Diversity, 9(1), 9. 10.3390/d9010009

12. Donald, F., Purse, B. V., & Green, S. (2021). Investigating the Role of Restoration Plantings in Introducing Disease—A Case Study Using Phytophthora. Forests, 12(6), 764. 10.3390/f12060764

13. Earl, D. A., & vonHoldt, B. M. (2012). STRUCTURE HARVESTER: a website and program for visualizing STRUCTURE output and implementing the Evanno method. Conservation Genetics Resources, 4(2), 359–361. 10.1007/s12686-011-9548-7

14. Ennos, R., Worrell, R., & Malcolm, D. (1998). The genetic management of native species in Scotland. Forestry, 71(1), 1–1. 10.1093/forestry/71.1.1-a

15. Ennos, R. A. (2015). Resilience of forests to pathogens: an evolutionary ecology perspective. Forestry, 88(1), 41–52. 10.1093/forestry/cpu048

16. Evanno, G., Regnaut, S., & Goudet, J. (2005). Detecting the number of clusters of individuals using the software STRUCTURE: a simulation study. Molecular Ecology, 14(8), 2611–2620. 10.1111/j.1365-294x.2005.02553.x

17. Fady, B., Cottrell, J., Ackzell, L., Alía, R., Muys, B., Prada, A., & González-Martínez, S. C. (2016). Forests and global change: what can genetics contribute to the major forest management and policy challenges of the twenty-first century? Regional Environmental Change, 16(4), 927–939. 10.1007/s10113-015-0843-9

18. Finger, A., Rao, S., Cowie, N., MacDonell, T., Beck, A., & Denny, B. (2022). Conservation genetics of montane willow populations in Scotland - limited natural recovery despite long-distance gene flow and high genetic diversity. Environmental Research: Ecology. 10.1088/2752-664X/ac9682

19. García, C., Guichoux, E., & Hampe, A. (2018). A comparative analysis between SNPs and SSRs to investigate genetic variation in a juniper species (Juniperus phoenicea ssp. turbinata). Tree genetics & genomes, 14(6), 87–87. 10.1007/s11295-018-1301-x

20. García, D. (2001). Effects of seed dispersal on Juniperus communis recruitment on a Mediterranean mountain. Journal of Vegetation Science, 12(6), 839–848. 10.2307/3236872

21. Garcıá, D., Zamora, R., Hódar, J. A., & Gómez, J. M. (1999). Age structure of Juniperus communis L. in the Iberian peninsula: Conservation of remnant populations in Mediterranean mountains. Biological Conservation, 87(2), 215–220. 10.1016/s0006-3207(98)00059-7

22. Green, S., Elliot, M., Armstrong, A., & Hendry, S. J. (2015). Phytophthora austrocedrae emerges as a serious threat to juniper (Juniperus communis) in Britain. Plant pathology, 64(2), 456–466. 10.1111/ppa.12253

23. Green, S., James, E. R., Clark, D., Clarke, T.-K., & Riddell, C. E. (2020). Evidence for natural resistance in Juniperus communis to Phytophthora austrocedri. Journal of Plant Pathology. 10.1007/s42161-020-00693-1

24. Gruwez, R., De Frenne, P., De Schrijver, A., Leroux, O., Vangansbeke, P., & Verheyen, K. (2014). Negative effects of temperature and atmospheric depositions on the seed viability of common juniper (Juniperus communis). Ann Bot, 113(3), 489–500. 10.1093/aob/mct272

25. Gruwez, R., De Frenne, P., De Schrijver, A., Vangansbeke, P., & Verheyen, K. (2016). Climate warming and atmospheric deposition affect seed viability of common juniper (Juniperus communis) via their impact on the nutrient status of the plant. Ecological Research, 32(2), 135–144. 10.1007/s11284-016-1422-3

26. Gruwez, R., De Frenne, P., Vander Mijnsbrugge, K., Vangansbeke, P., & Verheyen, K. (2016). Increased temperatures negatively affect Juniperus communis seeds: evidence from transplant experiments along a latitudinal gradient. Plant Biol (Stuttg), 18(3), 417–422. 10.1111/plb.12407

27. Hall, S. A. (1990). Pollen deposition and vegetation in the southern Rocky Mountains and southwest Plains, USA. Grana, 29(1), 47–61. 10.1080/00173139009429976

28. Hantemirova, E. V., Berkutenko, A. N., & Semerikov, V. L. (2012). Systematics and gene geography of Juniperus communis L. inferred from isoenzyme data. Russian journal of genetics, 48(9), 920–926. 10.1134/S1022795412090050

29. Hantemirova, E. V., Heinze, B., Knyazeva, S. G., Musaev, A. M., Lascoux, M., & Semerikov, V. L. (2017). A new Eurasian phylogeographical paradigm? Limited contribution of southern populations to the recolonization of high latitude populations in Juniperus communis L. (Cupressaceae). Journal of biogeography, 44(2), 271–282. 10.1111/jbi.12867

30. Hijmans, R. (2023). raster: Geographic Data Analysis and Modeling. In (Version 3.6-26) [R Package]. CRAN. https://CRAN.R-project.org/package=raster

31. Hubert, J., & Cottrell, J. (2014). Establishing and managing gene conservation units [UKFS Practice Note](FCPN021). https://cdn.forestresearch.gov.uk/2014/03/fcpn021.pdf

32. Klimko, M., Boratyńska, K., Montserrat, J. M., Didukh, Y., Romo, A., Gómez, D., Kluza-Wieloch, M., Marcysiak, K., & Boratyński, A. (2007). Morphological variation of Juniperus oxycedrus subsp. oxycedrus (Cupressaceae) in the Mediterranean region. Flora - Morphology, Distribution, Functional Ecology of Plants, 202(2), 133–147. 10.1016/j.flora.2006.03.006

33. Knyazeva, S. G., & Hantemirova, E. V. (2020). Comparative Analysis of Genetic and Morpho- Anatomical Variability of Common Juniper (Juniperus communis L.). Russian journal of genetics, 56(1), 48–58. 10.1134/S102279542001007X

34. Kopelman, N. M., Mayzel, J., Jakobsson, M., Rosenberg, N. A., & Mayrose, I. (2015). CLUMPAK: a program for identifying clustering modes and packaging population structure inferences across K. Molecular Ecology Resources, 15(5), 1179–1191. 10.1111/1755-0998.12387

35. Lefèvre, F., Koskela, J., Hubert, J., Kraigher, H., Longauer, R., Olrik, D. C., Schüler, S., Bozzano, M., Alizoti, P., Bakys, R., Baldwin, C., Ballian, D., Black-Samuelsson, S., Bednarova, D., Bordács, S., Collin, E., de Cuyper, B., de Vries, S. M. G., Eysteinsson, T.,…Zariŋa, I. (2013). Dynamic conservation of forest genetic resources in 33 European countries. Conservation Biology, 27(2), 373–384. 10.1111/j.1523-1739.2012.01961.x

36. Lowe, A. J., Cavers, S., Boshier, D., Breed, M. F., & Hollingsworth, P. M. (2015). The resilience of forest fragmentation genetics—no longer a paradox—we were just looking in the wrong place. Heredity, 115(2), 97–99. 10.1038/hdy.2015.40

37. McBride, A. (2005). Manging uplands for juniper (Back from theBrink Management Series., Issue.

38. Meirmans, P. G. (2012). The trouble with isolation by distance. Molecular Ecology, 21(12), 2839–2846. 10.1111/j.1365-294x.2012.05578.x

39. Merwe, M. V., Winfield, M. O., Arnold, G. M., & Parker, J. S. (2000). Spatial and temporal aspects of the genetic structure of Juniperus communis populations. Molecular Ecology, 9(4), 379–386. 10.1046/j.1365-294x.2000.00868.x

40. Michalczyk, I. M., Lücke, Y. A., Huck, S., & Ziegenhagen, B. (2008). Genetic support for recurrent fragmentation and founder events of juniper populations in Central Europe. In Application of DNA marker systems to test for genetic imprints of habitat fragmentation in Juniperus communis L. on different spatial and temporal scales: Philipps-University of Marburg.

41. Michalczyk, I. M., Opgenoorth, L., Luecke, Y., Huck, S., & Ziegenhagen, B. (2010). Genetic support for perglacial survival of Juniperus communis L. in Central Europe. The Holocene, 20(6), 887–894. 10.1177/0959683610365943

42. Oostermeijer, J. G. B., & De Knegt, B. (2004). Genetic population structure of the wind-pollinated, dioecious shrub Juniperus communis in fragmented Dutch heathlands. Plant species biology, 19(3), 175–184. 10.1111/j.1442-1984.2004.00113.x

43. Peakall, R., & Smouse, P. E. (2012). GenAlEx 6.5: genetic analysis in Excel. Population genetic software for teaching and research-an update. Bioinformatics, 28(19), 2537–2539. 10.1093/bioinformatics/bts460

44. Pers-Kamczyc, E., Mąderek, E., & Kamczyc, J. (2022). Seed Quantity or Quality?-Reproductive Responses of Females of Two Dioecious Woody Species to Long-Term Fertilisation. International Journal of Molecular Sciences, 23(6). 10.3390/ijms23063187

45. Pers-Kamczyc, E., Tyrała-Wierucka, Ż., Rabska, M., Wrońska-Pilarek, D., & Kamczyc, J. (2020). The higher availability of nutrients increases the production but decreases the quality of pollen grains in Juniperus communis L. Journal of Plant Physiology, 248, 153156–153156. 10.1016/j.jplph.2020.153156

46. Pritchard, J. K., Stephens, M., & Donnelly, P. (2000). Inference of population structure using multilocus genotype data. Genetics, 155(2), 945–959. 10.1093/genetics/155.2.945.

47. Provan, J., Beatty, G. E., Hunter, A. M., McDonald, R. A., McLaughlin, E., Preston, S. J., & Wilson, S. (2008). Restricted gene flow in fragmented populations of a wind-pollinated tree. Conservation genetics (Print), 9(6), 1521–1532. 10.1007/s10592-007-9484-y

48. Reim, S., Lochschmidt, F., Proft, A., Tröber, U., & Wolf, H. (2016). Genetic structure and diversity in Juniperus communis populations in Saxony, Germany. Biodiversity research and conservation, 42(1), 9–18. 10.1515/biorc-2016-0008

49. Reynolds, F. (2022). The current status of Juniper (Juniperus communis) in Wales: an assessment of site and genetic management Aberystwyth University].

50. Robledo-Arnuncio, J. J., Collada, C., Alía, R., & Gil, L. (2005). Genetic structure of montane isolates of Pinus sylvestris L. in a Mediterranean refugial area. Journal of biogeography, 32(4), 595–605. 10.1111/j.1365-2699.2004.01196.x

51. Rodriguez, A. C. (2019). Quantitative Genetics and Genomic Selection of Scots pine Swedish University of Agricultural Sciences]. Umea.

52. Salmela, M. J. (2011). Adaptive genetic variation in Scots pine (Pinus sylvestris L.) in Scotland University of Edinburgh]. Edinburgh.

53. Salmela, M. J., Cavers, S., Cottrell, J. E., Iason, G. R., & Ennos, R. A. (2013). Spring phenology shows genetic variation among and within populations in seedlings of Scots pine (Pinus sylvestrisL.) in the Scottish Highlands. Plant Ecology & Diversity, 6(3-4), 523–536. 10.1080/17550874.2013.795627

54. Stace, C. (2019). New Flora of the British Isles (Fourth ed.). C & M Floristics.

55. Sullivan, G. (2003). Extent and condition of juniper scrub in Scotland (026). (Archive Reports, Issue. https://media.nature.scot/record/~8e94e99258

56. Sullivan, G. B. (2001). Prostrate Juniper Heath in North-west Scotland: Historical, Ecological, and Taxonomic Issues University of Aberdeen]. https://ethos.bl.uk/OrderDetails.do?uin=uk.bl.ethos.369544

57. Surso, M. (2018). Pollination and pollen germination in common juniper (Juniperus communis: Cupressaceae). Arctic Environmental Research, 18(4), 162–174. 10.3897/issn2541-8416.2018.18.4.162

58. Team, R. C. (2021). R: A language and environment for statistical computing. In https://www.R-project.org/

59. Thomas, P. A., El-Barghathi, M., & Polwart, A. (2007). Biological Flora of the British Isles: Juniperus communis L. Journal of Ecology, 95(6), 1404–1440. 10.1111/j.1365-2745.2007.01308.x

60. Unterholzner, L., Carrer, M., Bär, A., Beikircher, B., Dämon, B., Losso, A., Prendin, A. L., & Mayr, S. (2020). Juniperus communis populations exhibit low variability in hydraulic safety and efficiency. Tree Physiology, 40(12), 1668–1679. 10.1093/treephys/tpaa103

61. Van Oosterhout, C., Hutchinson, W. F., Wills, D. P. M., & Shipley, P. (2004). microchecker: software for identifying and correcting genotyping errors in microsatellite data. Molecular ecology notes, 4(3), 535–538. 10.1111/j.1471-8286.2004.00684.x

62. Vanden-Broeck, A., Gruwez, R., Cox, K., Adriaenssens, S., Michalczyk, I. M., & Verheyen, K. (2011). Genetic structure and seed-mediated dispersal rates of an endangered shrub in a fragmented landscape: a case study for Juniperus communis in northwestern Europe. BMC Genetics, 12, 73–73. 10.1186/1471-2156-12-73

63. Verheyen, K., Adriaenssens, S., Gruwez, R., Michalczyk, I. M., Ward, L. K., Rosseel, Y., Van den Broeck, A., & García, D. (2009). Juniperus communis: victim of the combined action of climate warming and nitrogen deposition? Plant Biology, 11 Suppl 1, 49–59. 10.1111/j.1438-8677.2009.00214.x

64. Verheyen, K., Schreurs, K., Vanholen, B., & Hermy, M. (2005). Intensive management fails to promote recruitment in the last large population of Juniperus communis (L.) in Flanders (Belgium). Biological Conservation, 124(1), 113–121. 10.1016/j.biocon.2005.01.018

65. Ward, L. K. (1982). The Conservation of Juniper: Longevity and Old Age. The Journal of Applied Ecology, 19(3), 917. 10.2307/2403293

66. Ward, L. K. (2007). Lifetime sexual dimorphism in Juniperus communis var. communis. Plant species biology, 22(1), 11–21. 10.1111/j.1442-1984.2007.00171.x

67. Wickham, H., François, R., Henry, L., Müller, K., & Vaughan, D. (2023). dplyr: A Grammar of Data Manipulation. In (Version 1.1.4) [R Package]. CRAN. https://CRAN.R-project.org/package=dplyr

68. Young, A., Boyle, T., & Brown, T. (1996). The population genetic consequences of habitat fragmentation for plants. Trends in Ecology & Evolution, 11(10), 413–418. 10.1016/0169-5347(96)10045-8

