## Appendix for "Evidence of genetic isolation and differentiation among historically fragmented British populations of common juniper, *Juniperus communis* L."

Appendix 1: The 74 post-filtering loci returned from the two multiplex Sequenom SNP chips from the CNRGV, including loci names, the two SNP base pair genotypes, which multiplex chip the loci was tested in, and the full sequence, with the SNP site and genotypes delineated with square brackets.

| Loci | SNP genotypes | Multilplex 1 or 2 | Sequence |
| --- | --- | --- | --- |
| 1082_62 | A/G | 1 | CTGCAGGTTTAGGACCCCTTTTGCCGCCCTTTGGAGGAGCCTTCTCCTTGGGCTTCTTCTC[A/G]GCAGGGCTTTTCTCTCCAACCGTCTTCTTCTCTTCAACCTTCTCCGCAGGTTTCTTCTCTGCGGGCTTCTTCTCTGCCTTGGGCGCCATTTTTCTGATCAATAGCAATCAATAAGGAAAATAAAAACGAAATCCTTAT |
| 10210_101 | T/A | 2 | CTGCAGCATTATCAGAGTCTGGGGTATTTCAGATTCTATGGATGTGTATTGCAGCAGGGGCAGTGAGTTATGTGGTGGCATTCCAGAGCAGATGCGCCCT[T/A]CCTGGACCTTGATCAATCTCATGGCCAGAGAAATATTGACAACTGTGGCACAGCCACCCCACTATGCATAGGGGTTATCCACAGTGGCTAGAGCATATA |
| 11457_34 | A/T | 2 | CTGCAGACCATCATGTTTCAGAATTTGCTTCAT[A/T]TACCAGTAAGACATCCTCGCAGATTTATTTCTTTCAGTTTTCCTTCACTCTGATGAAATAGCACTTTGATTGCATTTGGAGAATATAAAGGAATTTATCAACTTTGTTTTACCTTAATGAAATTTCATGATGCCGGCAGATTAGTGCTGGAAGCCCATTCAGATAC |
| 11742_169 | T/G | 1 | CTGCAGCAAAACTTCTCATCTTGCCGATCATCCTGTACGCCGACAGCGTCTTTGTCTGCTGCGTGAAACACAATGCATACGCTGGACCACCATTCACGTTCATTACTGTTTCAAACACCACAAGATGCACCAAGATCTGATTATAATTATCTAAGAATAATGCCATAA[T/G]CACAGTGGCCAAGTTCATCAGAATCATGAAA |
| 1198_43 | C/T | 2 | CTGCAGCTGAATAGTCTTCCTCTAATATCATTTCTTCATCTT[C/T]GAAGAAATTAAATTCAAGAACAATCTGAATAGGTTCATTCTCTTTAATGAAGTTATGCTTACCAACTTAGCCTCGTGGCTGCAAGCAGCATTCCCATGTTCATTTCTATAGCCTGCATGACTTTGCATGTAACCTCTCATCCTCATGTTGGTTCTTG |
| 11995_176 | G/A | 2 | TGCAGAAAATATCAATCTTCCTCCGGCACTGCTTTCCCGTTTTGACTTAATGTGGCTGATACTTGACAAAGCTGACATGGATAGCGACTTGGAAATGGCTAGACACGTCTTATATGTGCACCAAAAGTGCGAGCCACCTCCTTTGGGTTTTGCTCCTCTTGATTCATCCATACT[G/A]AGGTGAGTAATCGACTGATCCTTA |
| 1212_81 | C/T | 1 | CTGCAGAGAAAATCAGTATTGCTCTGCATAATCCTCCAACATACACATGTAGTTGAGAAGCATTCTCTCATAACCTCCCT[C/T]CATCATTGCATCTTCATCTCACCTGGGGAGAACAGCATTATCTTTTTTGCATCTCAATCACTCACATGCACAACACTACACATTTGGCTCATCCTACTCTTGCATACACAATCAGTCTT |
| 12138_151 | C/T | 1 | CTGCAGATTTGGTACTATCAGATCCTTGTTTTTCTTGCTGGCACATTGAATGATCCAGAAGTTGATATAGATGCCCTTTCAATAAGGTAGTTAATCCTGATGCTTCAGGCCATTGCTTAAGAAAGGTGATTAAATATGGATATATTTATA[C/T]TTGTCTAATTTACATTTTGTTGATAATTTCAGCTATACAGAACAGTTGA |
| 12196_162 | C/T | 2 | CTGCAGAATTGACAATGATCTTAATCTGAAATTAAAGGAAGTAAATGAATCCTACCTTAAATCAAAATAATTTCCTCGAAGACAAAGTATAAGTCCAATGACACCAATGATCAAAAAGGGGAAGTTTGATATCACATTCAGAGTATTGGGCACGCCTATAA[C/T]ACAAACACAAATACTTGTCAATATATCTCCACAAAATG |
| 12684_84 | G/T | 2 | CTGCAGCTTGGCACCACATAGTCAAATACATTGGTTGACTTGAGTAGTATCTCAATCAGCTCAATGACATTCAAATTGACTCA[G/T]CACTCTAGTTGCACATTGCCTTGTACACACACCTTCCAACGCAAATACTAATGATTGGAAGACCAATGGTATTTTAATTTATATTCTATCACTTATTGACAATGTGTAATTTTTTT |
| 13131_128 | T/C | 1 | CTGCAGCATTTGTGATAAAAGGGGGTTTATTGGCTACTGTAAAGGTCACTGGTACACACACACTACATATGCCTATTTATGAATGTATGTAGATGATTTTGTACTTATTTTGATACCGTGCTACTTA[T/C]TGTAATCAGCTTTATGAAATTTTTCAATGTTGATATTCATGTTGCACTTTTTTTATGTTCAAGTATTTGGTA |
| 13205_45 | C/T | 1 | CTGCAGTGGGGAAAAGAAAAGTCACGCCTGAACAAACAAGTCGG[C/T]GAAGGAGTTTGAGCCTGAGGTCGCTAAAGAAAACATAGCCCGTGGCTCACATAAGGTGCTTGTAGACAACACAAAGTCCTACAGACGATATAAGAAGCGGAAAGGGGAAACAAATTTCCTGTGGACATTGCAAGATGAGACGGGCTGTCTAACCT |
| 13997_141 | A/G | 1 | CTGCAGATTTGGATCGTAATCTTTTTATAGTCTTCGAATATCTTTCTAATTGTGATGGTATAGCAATTTAGCCTTCGACTTGGTCATATATGTGTGCTAATACTTCATCGCTCCACAGTATTTATTGAATAATTTAATAT[A/G]GTCTTCAACTCATGCTCTGTGTTGTTTAGATGGCTCTGTGTTGTTAATAAATTGGTACT |
| 14296_68 | T/G | 1 | CTGCAGTCATCAGGGTCCCATATTTTCATTGAAAACAAAGGACAACACATAATATATGGCAACACAA[T/G]AGTATTTCATAGTAAAAAAATTACCAAATTGGTTTGGAATGTAAATTATTTTTCAGGTTTATCAGTAATACATCGTCAATTCCTCTATGGCATAAGCAACCATTTGCCTAAACCATGAAAAGAAGAGAAGAA |
| 15494_81 | C/T | 2 | CTGCAGAGAAAATCAGTATTGCTCTGCATAATCCTCCAACATACACATGTAGTTGAGAAGCATTCTCTCATAACCTCCCT[C/T]CATCATTGCATCTTCATCTCACCTGGGGAGAACAGCATTATCTTTTTGCATCTCAATCACTCACATGCACAACACTACACATTTGGCTCATCCTACTCTTGCATACACAATCAGTCTTT |
| 1641_85 | C/T | 1 | CTGCAGGTTTCTGTGAAATCTAATTGGATTTTTTCCGCAGTTTGTTCTATTGACAAATCAATTTTTGGTTCAGATTTGGTATTA[C/T]TTCAGTCTGTTCATGTTGTATTTTAAAATCTTCATCTTTCCAGATAGCTTCTAAATAGGCCAATTCTTCTTCTTGATGAGGTAATGTAGGATTGTCCTAATTTTCAATTTCATTT |
| 16700_165 | T/C | 2 | CTGCAGATCTCTCAACTGATTCAGAGAATAATATCTGCAATGTGAACTTTTTATTTAACTAATGTTTTGGTTACTTACCATGCAGGAAGTTTTCCTTACAGTTTGGCAGGTAGCTATTATGTATTATCCGGCTGCTTATTTTAATGAATATCTGATTGCTACTT[T/C]TCTTTTCTTACTTCTCTATGTCTGCTGAGTGAGGC |
| 16751_50 | A/G | 2 | CTGCAGACCAACAGTTGTAATATAATGGTACTGGGGATGTAATATATTT[A/G]GGATTCCTAGGAGTGTAACACAGCTTCATCATCACATTGCAATAAGGTGGCTCCTATTCCCAGTATGTAGCAAGAATTTGTACAACTAGCACAAAAATGTGGATATATCAAAACCAAAAGAGAGACAATTATAGAAGAAAACCCTAAAAT |
| 16960_53 | A/C | 1 | CTGCAGGCAGCTATTGAAAACAAAGTCACTGAAGTCTGACGCTCTGACAATG[A/C]ATTTATTTCATAGTTATCTTATCCTTTCTGTCAAAAGTATTTGGTTTTAAAAATATTTAATGGATAGGCATCTTGGCCAGACCAAAGAACAATCGATCAGGTAACCGGTTCAGACTTTATCATCCAGTGGCTGTAGTGCTGTTGAAC |
| 17261_133 | A/C | 2 | CTGCAGAGTTTGTGAGAAGATATTTATTATCTGATCGAAGTTTCTGTTAGCTTTCATCTGAAAATGTCAGTCCATGAATTTGTAGATTGGGATACAGATGACTTTGATGGGGTAACCGAATTCACCTGGTGC[A/C]CAAGTACTGATTTCATCTGTGTTGTGGGCTATATTTATATAACCTAATAAGTAACTTTTAGTAGCAA |
| 17598_91 | A/C | 2 | CTGCAGGTCACGATCTTGGAACCTATCAGGAACCAAAAAATTCTTTGAATCATTTTTTAAAGTCTGATATTTAAAATAAAGACTCGCAGA[A/C]GCATGAGTCCTCATAAATAACATTCGTAGTACCCAAATCACAAGTACCAACATCATCCACCTATAAAACCTCTGCTATCTCATGTAGATGTGAACATAACTCACTGCTG |
| 17630_62 | C/T | 1 | CTGCAGAGCTGGAATCACACTTGGCCAACAGGAAGGATAAAACAGATAGTGTTCTTTAAAT[C/T]GGCTTTAATGGATAAAATCAATCCTTTTTGCCACAGCTTGCTTTCTCATAGTGCCTATTGTTATCTCTGCATACATACTTAGATCTATGCTGCATTTGTTGCTTTTAGATTTAGTGAAGTCAAGATGTAAGACCTCGT |
| 17676_127 | A/T | 2 | CTGCAGTTGTTTATAGTTGCAAAACTGCATGTAAGGGTAAGCTCCCTTTGAGTGGAGCTTCCTAAAATAATTTGGTGGTTGCTGCTGAGAATTTTTAGGAATATTCTGAAGGTGATCGTCCTATTA[A/T]GTCAGTGGTGCAGCTCCTTAAAAATTTAAGCTCTGAACTTTAGGGTAGATGGCGGTCTCATGAAGATTGCAGC |
| 18371_155 | T/A | 2 | CTGCAGCCTGCCCTAAAACGTGGTGACATTTTGAAAAAGAAAATGCTAATTTCAAAATTCAGCACTCTGTAGATAATTTTTAAAATTTCAGCTTCTATAAGTGTTCAAGTTATTTTAATAGCTAGCTCCCTGTTGTGCCCAAAATTCGTCGAAA[T/A]TTTTTGTTATCCAAAAACCAAATCTCCATATCCAAAAACTCACAG |
| 18536_64 | T/A | 2 | CTGCAGTCACTTCCCTCATTGAGCACACATACACATCAATAGTTTATAGACAATACAAAAAAT[T/A]AAGACTCGGTTGAAAATAATATTAACAAAACAAAAGGCCAGAAGAGATCTCACATATCCACACTAAATATCAGCCAAGTACCTGTAAGCTAGTTAATACTCTTGCTTTAAGGGTGTGATCCATGGTTAGCTCATGG |
| 18825_68 | A/G | 1 | CTGCAGATGAAAGCCATCCACAAATTCAATTGCTAACTTGCCTGCAAAACTTGTAATCCCTTGCCAA[A/G]ATTCCTTTTTTGAGATCCAAGAGCAAGGAGAGTCCTAAGATACTCGAAGGGAAGAGTTCCTATTTGAAATCTAAGAATCTGAGGAATCTTGTTCTAGATAAGCTGAGGGGTATTAAATGAAAAGATAGAGGA |
| 18882_160 | A/G | 1 | CTGCAGGGCAAAATACTTTAGCAATGCCTTGTAGACCTCTATACCTGCATACATATCCTTGGCTTCCATCTCTCTAATCAAAGACTCTCCTGCTTCAGGCATTCCTACCCTCACATAAGCCATTATCATTGCACCATATGCTTTTTGATCCAGTTGAAG[A/G]CCAGATAGTCTCATCTTATCAAATGTATGTCTTGCCTTAT |
| 19160_119 | T/G | 2 | CTGCAGGGTGGGGGGCGCCCTTACTTATAATAGTAGGGGGTTAATAATAGTTGGGCAACTATAATCCCCTTCATGGCCCTACCTAAAATTCTATGGATGGGCTATGGGCCTGCTAGAG[T/G]TGCCATTACTATGCCCCTGCACCACCCCCCCAATAATAAGTTGTTGAATAGTTAAGATACCTCCAACTATAATGAGAGGAG |
| 19165_137 | C/G | 1 | CTGCAGCTCTATCAGTTCCATATTTTCAAGACTGACCTAACAAGTAAAAGAAGAATGTGGGTGCTTCTTTTGTTTAAAACACATAAGATCTGTGCAGCTCTATTAGAGCCATATTTTCAAGATTGACCTTGCACAC[C/G]TTTACGACAAATCCTTCAAATTTAAAAATATCAAAACACATGATGGCAAGAATGGATGCAGGC |
| 19553_85 | G/A | 2 | CTGCAGGACACATGTCAGGGTCCTATCTCTGCCAAATTTCAAAGGGCTGAAGCTAAAGCCTTGCAATATAAACGTCTTTTTCAA[G/A]GAAAGTTATGGGGATTTGTGTTAAAGGAAAATGTTACCACACAGCAGAAGGCAGGCATTCGCTGTGGAAGGAATTAGTTAACTTTTCATGTGATGGAAAATCATTAAGAAAAAGT |
| 19809_45 | G/A | 1 | CTGCAGTCTATATTTATTCCTCTGGTAGGTGACAATAGTTATAC[G/A]TTCAAATTCAAATTCACATTCAACACTACTTTCAACATGATACATATTGCAAACTACAATGGTTATTCCAATCTATTCCCTTTCACCAAACTGTGCATAATTAGTCTCTTGGAAATGGACATTGGAGGGAAATTTGAAAAGCCAAAGTTCTTGTG |
| 19942_106 | T/A | 2 | CTGCAGATGATGTCTATCTGACTGTTGGCTGTCTACAAAGTATTGATTTTGCTATGAAAGTCTTAGGTCGTGAAGGTGCAAACATTCTTCTTCCAAGGCCAGGAT[T/A]TCCATATTATGACTCAATAGTGGCATACAAAGGTATTGAAGTCAGGTATTATGATTACTGCCAGAAAGAGATTGGGAAATTGACTTGGGTCAAG |
| 20726_122 | C/T | 1 | CTGCAGCATGATGTGCAGGAGCTGAACCGTGTGTTGTGTGAGAAACTAGAGGACAAAATGAAGGTCTTGAATTTTGTATTTAATCTGTGATTGTCCCAGCACCCCAATTTCTCATGAAGTT[C/T]TTGTTGAAATTGAAAGCTTCTTTATACTGTTATATTTCAGGGTACTGCTGTGGAGGGTACAATACAACAACTATTTGA |
| 211_106 | A/T | 2 | CTGCAGCATTGCATGTAAATAGGTGCATAGTAAAGCTTTTAGGAAAAGGTCAAGAAGATGTTCTAAAATGAGACTGAAAAACTACATAGAAAAAGGGTACCTGCG[A/T]GCTAATATGTCATTTTACTGTACAGAATGCGGGTCACTGATGGAGCTGGACTATCTGAGAAAGTTGACAGTTCATCAATCTAAATTCAACAAAA |
| 21351_162 | G/A | 1 | CTGCAGCCTAAGAAGAGGAAATCCAATGATAACAATGAAACCCAACTGTGTAGGGGATCCAGATGCACAAAATCAACTCATTTCCAATTTTTCACATATCAGATTTTCTAAGAAGAAACATTTTCACCTATCAAATCTTCTTCTGAGGATGAAGAAACACA[G/A]AACTATTCTATCTCATGTCGATTTCTCTGTCATGAATC |
| 21440_152 | T/C | 1 | CTGCAGGCATAATATACAGAATCCAAGAAAACTTTATAAGAAAAAGAATGGATGTTATCATGTTCTGTCTTATGGCAATTGATATGTTATAACAGCTAATCATGTCAACCACAAATCGCAAAGTTTATAATTTACTAGTGTCAGTAGATAA[T/C]GATTTCATAATAGGGAACAAATCAGTGGAGAAAGGGAAACATGTTCAG |
| 21548_155 | C/G | 1 | CTGCAGATGATCGCAGATTAGAGTACTTAATATAGAAGTATGCAGATTGCAGTAACTGGAGATAATAACATGTATTGTGGTGGATGCTTTCTCTCAATCACTGGATCAATATATATATATGAGAACAAAGGGTCATGTTAATGTAGAGACATGA[C/G]CCTACATGGCATGAGCCACATAGGGTCATGTTCCTACATCAACAT |
| 21832_135 | T/C | 1 | CTGCAGTGCAAGAAAACACCATGAGAAATAAACATTTTCAAAAATATTTATCGTTCTTTTCCCAAAATAGAAAACTATCTATGTGCTTTCAGGTGCTCTGATGTGCTACTCTCTAGGTGATTTATTTGCAACCA[T/C]ATGCTGACATATAAGAAATTTTGGTCCTTACATACAAAATATGATGTCAACATTCAGCTTCAAGC |
| 22176_62 | C/T | 1 | CTGCAGCTCGAGGGGACGGATCCATGGTGAGATTGCTGATGCAGAAAGGGGCTACGAAAGA[C/T]ATCAGGAACAAGCACGGGAAGACCTCATATGACCTTGCGGTGGACACAGGGGACTCCGCTTTGCTCGATTGCTGCGTCTCGGCGATGGCCTCCGTAGGGCTGCAAGGCGCGGGGATTTGCGAAGTCTGCAACGGTGCC |
| 22479_130 | T/C | 2 | CTGCAGAATGTCTACTACAGTTGGAGCCACAGAATGCTGGTATACGTGTGTTGCTTTCAAACATATATGCTGAAGCTGGCAGGTGGGATGATGTAAGAAAAGTGCGAAAAATGATGAAAGACACAAAGG[T/C]TGTAAAGGACCAGGACAAAGCTGGATTGAGATCAAGGAATCAGTGCATGTGTTCTATGCCGAAGACCAAT |
| 22617_130 | C/A | 2 | CTGCAGTCGATCTGCCTTGCCCACAAAAATGGCTGGTAGGATATGTAAAACAGCACCATAATGGCTTGTCGATCTAGCAATCTCAAATTGTGATCGGAAATCAATATCAACAATTAGCCTCTCCGACTT[C/A]CCCATCGGTCCCACTCCTTCCAGGATAACATCAATGTATTCATACTCTCCTGAAGGAAAATACAAATGTA |
| 22877_36 | G/A | 2 | CTGCAGATTGGGGTGGAACAGAAAAATGGTGATGT[G/A]CAGGAAACTAAATGGAGTATAAGTTCAAACCCTAACAGCCTATTCGTTGATGTAAATGAAAATAGTTCTCCCTGTACTCTGATACCAATGTCAGGCCAGGAAATTGGACTGCAAAGCAAAATTTGACATTGTATTGATGCAAAATCCAAGATGGGATTATTGTT |
| 23028_86 | G/T | 1 | CTGCAGTAATGGACAAAGACGCTATAGTCTTTCACTTATTCATTTAACTGGTTGCACATAAAATAAAGTAAATGCATAAGCTTTG[G/T]TGGATCTTTTGTTATCAATATCAATATCCCTATTTGCATGGTTTAATAATAAAGGATAATAAATTTAAAATACAAAAGAGTAATAATAAAATTTAAAATTAAAAGGGTAATATT |
| 23735_62 | T/G | 2 | CTGCAGATAATAAAAATACATATACACTGGGCATATATTTATAAACTCAACAACATAATAG[T/G]TTGAAAGATAGAGTTAACAGAAATCACAAATTAAAAGGTGTTATATATTTAAGACTTAGGGCAATATTAAGCAAACTGGTCAAACACAATAAATGGAGCAGTGATATGTATGATGAATCTATTAAAATAAGAAGGCCT |
| 23845_33 | T/C | 2 | CTGCAGTAGAAAGGAAAATCAGCACACCAATT[T/C]CAAAAGTATTTTAAGTGCAGATAAATATTCATATACATATTGATGTCTACACTAATGCCCACTTTTTATCTATATATTTCTTTCTTACAGAAAACGTAACCTAAACAATAATTTGATTGCAAATACTAAAATGGTAATAGCACCTAGGAGAACATAACAACACATGA |
| 2390_145 | C/T | 2 | CTGCAGACACTGAATGGCCATGGGATTGAGACCGGAAAGGGCTTGGCGTGCAAACTCGTCGTCTGTCTGCCACGCATGCTCGTCAGCTGCAAGACAGAAATAAAAGTTCAAATTTTATTGTACCAATGATAGGTTATTCGATAA[C/T]TAGTACTAGCATAAAACCGCTTGAAATTTATTTAAATGAATGGTGGTGATCTCGC |
| 2715_44 | A/C | 2 | CTGCAGTGTGAGTGAATGTTTTTAACCATCACCAACATACTAG[A/C]CAGATTATCATTGATAAGTTTTTCTTTTGTTTTAAACTATGTTTAAAATTTTTGGTCGATCTGTCTAAATTGTGTATTATTATCATGACGATTGTAATGTATGCCACTTAATTTTAAACGCCTACTTTTCATATGCTTTAAAAATATCACATTAAG |
| 2815_174 | G/A | 2 | CTGCAGTGCTTTTTCTGCAACATTTGCCCTTTTCGTTTCAATTGCAGTTCTACTGGTATCCCCTGTTTCTGCTGCCCTAATTTTCTCCCGCGCCCTGCTTCCCCACTCTTACATCTGACCGGATTTGCTCTCCAAATCCGTGAATTCGTCAATTCCGCCGGATTTCCGAATTC[G/A]CGCTACACGCATCTGATTTGCAGCAA |
| 3099_117 | T/A | 2 | CTGCAGTCATTAGAACCTATGCATTTCTGCAATAAGATACAAATCGCTAAAATAAGTGGCAATTGAAAAGCAGTTGAAAAATATGTTTTACTCTAGCTATTATGACGATCTGCAAT[T/A]TTACAGATATTGTGATGATATCACTGAAAGACAAAGAAAAATGGAATCCAGTAAAGGACATGCCTTTCATAGTTGAGGGCACA |
| 3293_161 | T/A | 2 | CTGCAGAACAAGTACTTTGAATATAGGCTCACATGCTCTATAATCCATGAAAGTCAGTATCGCAAGTGCCTGTATATAGGTAGAATTACGAAGTAAATGGGCTACTCTACGTGCTTTATTAGAAGACATACGTACATGCTTAGCTACAGCTCGTATTTTT[T/A]ATAGTTGAATTTAAATCTAAATCCCTATTTTGAAATATC |
| 3946_69 | A/G | 1 | CTGCAGCGCAGGCGATAGTCACTGAGCTTGGAATGTCCAATCCTAATGACAAGGGTGAAGTTGTAACA[A/G]AACACTTACTAACAACAATGAAACATCAAGTAGGTGGTCAGGACAATACATCATTGTTAACAACTATTCTGAAGAATGCTGGTATAATTGCAAAAGCCTTGGCCTCTTTGCTCTCTGGAATGGTCCAGGAC |
| 4003_165 | G/A | 1 | CTGCAGTTGGCACTAGGCTATCAATATTTTTGCTTGTATTAGAATACAAGCTTAATTTTTGAAAGATAGGGACCAGGCATGAATCTGTGGGATGAATGGCCCACGACAGAGGCTTTCACATTAAATTGTGTCAACCAAATCTAAAGCAATATTCTAAAAACCAT[G/A]CTCAACTCAAGCAAGGAAGTCATAAAAGAATCATT |
| 4131_172 | C/T | 1 | CTGCAGCTTTGGTACAAAACTAAAAGGCGAGAGATTTCTATTTCCTTTCATTGATAATTGATAAATCGTGATTAAATTTGTTTTTGATATGCGACCTATGGAAATTTAAGTCAATATAATAAACACTTTAATTTTTGAGAGAGTAATTACCTATGAAGGGATTGACTTTTT[C/T]TTAGTGTATGAATCATTTATCTATAACC |
| 4227_95 | A/C | 1 | CTGCAGAAATTTCGTAGAGCTAAAAAGGTATTAGACCTCGTTTATTTTTCCAATCTTTGAACGCGGGTAAATTGAGAATTTTCTTGTTCTTCCT[A/C]TTTTTCAATGCCCTCTGAAAACAAAACATGGCATGTTATGAAGAAATTAGCTGCAACTTTTTACCGCGCATGTACTTCTGAAACGAAGCAGCTCCAGTTCATTCC |
| 4497_57 | C/T | 2 | CTGCAGAGACCGTCAATAGCCTAAAGATTGTAAGAATAAATAGCATAAACCCTGCC[C/T]ACATTTCGTTTCTTTCCTTGCTGATCTATAACATGTTAGGTTTCACAATATTCTATTTAAACAATTTGAAATGCTGAGAGAAAAAGCCGAGAAGAATTAGCAGTTAGTTTGTGTTTCCCAGAAATTTGTTCTGTTCCTAAAAT |
| 465_128 | G/C | 2 | CTGCAGCCGGGGCTATATTCTTTAGCTTTTTGGTTAGCTTAGAACGGTTGCGAGTCATGATTACGTTGACAACGCGAGGCATGCTGGTGCATCGGAATAAACCCGGGCCCAGCAGTGGATGGGAAGG[G/C]CCACCCACTGCTTTATTGGCAATGCTGGGGTGGCCGGGTGGCGCACGAAATTTGAGCCGATTCTTCCGATCC |
| 4913_26 | C/A | 2 | CTGCAGATGTGAAATTATAGATACT[C/A]ACACGTTCACTTTGCCTAAGAATAATAATGCCACATTGATCTGTCTACATGCTCATACGGTCAATCTACTCTATGGATGTTATTGATCTGCTTATATTCCATTGGTACTATTACATGCACGACTGACTTACTGAATATAAAACAATCTGTTACAGGACAATTACAGTAAGATCA |
| 5134_143 | C/T | 1 | CTGCAGCATTCATGGGAACGTCTATGTCCAAAATACCCATGTGTTGTTTCAGAATGGCTCTGTGCTGAGTATTAAGAACTCACGTGATGGCTCCATCCAAAAAATGCCCCAATCGACTCCAATATGCTCATTCCTCCCTTTT[C/T]GTACTAGGGCATCAAGAAAGACATTCAACCCATGGACTTGTTTGTTTGAAGCTTTTT |
| 5447_151 | G/T | 2 | CTGCAGAAGCAGTAGGAAGAGGAAGGACCTCTCCCATTCCTTGCATTTATATGTTGTTGCTTTGATGCTTATGGTTGAAGTCAGAAAATTGTGTGAAAGTAGCTCTACCCAACTGCATTAGGCAAAGAGATCCAGTAACTGTTAGGATTA[G/T]GGGGGTAATGATCATGATAGAGTGTTGTCTTAGATACTTATAATAGCTG |
| 5662_100 | C/G | 2 | CTGCAGAGGACTCTCCTGCTAGTCAACGTAAGGATGGTCAGCTAAACAAATGAACCGGCTATAGCTATAAAAAATGCTTGAACTCAGCCTGAACTGCCC[C/G]GTAGGGCCGATGGGTGATTATGCAGAGGCCATAGAGATTATAGGCGATAGTTGTTTACTGTCCCCCTATATATTGAAGGGCTCGCCCTATATATTGAAGG |
| 5822_134 | G/A | 2 | CTGCAGAGTTATTTCCTCATTCAATCTTATCTTCTGCTCTTTATCCACTCCACTCTTTTTTTCTTACATATTCTCTGTATTCATTGTACTTCATTTTCTCATGAATGAATTCTCTCTTCCCTTTATATCATTC[G/A]TATTTTTCATGAACAATTGTGACTCTCTAAATAGACATCTCTTTTCAATATATATTACATCGCACC |
| 5868_118 | G/A | 2 | CTGCAGTCTGTTTGAGTCATTTACAGCCATCATCTTGCTGCCATTACACTGTACTAGAGCCATGATGTGTAGTGCAAGTTCCAATCAGTCCTGGGAGTAGCTAAATCAGGGTTCAGA[G/A]GCTCCACAAATGTTAGAGAAGCTCTGTAGCTGTTCTAGTGTTGCAACATCAATCATTATTTGGTATGTAAGAAATAAGACAC |
| 644_98 | G/A | 2 | CTGCAGGGCCATGATTGGCTAACAGACTGAAAGCGGGTGATATAGATGCAGAAATGCAACGTTCTTCTGGTTGCAACGATAGGCTTCCAGATGGATG[G/A]AAAAGTCCCACAATTGGATGGGATGGGATAGAAGAGGCGCCTAGTGGGGCTCCTTCTGATTCTGCCATAACGCATCTCTCCCCCCCTACTATTTTAAGGGCT |
| 6837_73 | G/T | 2 | CTGCAGGAAAATCATCCAAACAAAAAGGGGCTGTGTTCTATTTGGGTAAAATCATAGACAACCTAGATAAAA[G/T]GAGTATCTTTTAATTACTGGCATTTGCCATCTTTGAAATTTTGGCTTTGTGCTTATCTTTAAATGCTCCAAGATGGTAGATAGCTTGGTTGGACACTTAAATATATTGCTGTAATGTTGTCAATATA |
| 6912_133 | G/A | 1 | CTGCAGCATCATGCATGCTAAAGCGTTGCAGAATTTCGAAAGGCCTAAATGCAGGACAAAAGTTAGTTTCTACTTTATGCATTCCTACATAATTTCTTGGCAAATATGTAATCTTGAAGTATTAAGCAATCC[G/A]AATATAAGCCATGTTTGCGAATTTTACACAAGTGTACCAGTAGGAATATTAAAGAAAATAGGGGCTC |
| 7145_33 | T/C | 1 | CTGCAGTAGAAAGGAAAATCAGCACACCAATT[T/C]CAAAAGTATTTTTAAGTGCAGATAAATATTCATATACATATTGATGTCTACACTAATGCCCACTTTTTATCTATATATTTCTTTCTTACAGAAAACGTAACCTAAACAATAATTTGATTGCAAATACTAAAATGGTAATAGCACCTAGGAGAACATAACAACACATG |
| 7147_80 | C/T | 1 | CTGCAGATTCCCTTGGATCCACAAGTTATCAAGCACAGTGGATGGCACCACATTCGATATCCAACGATGAGGCCTTCCT[C/T]CCTATATATACTTTTCCTCCAAGAGGTTGGCCTCCTTCTAGAAGTCATAAAATAACATTTAATTTCAATTTCCAAAAGATTCACTTGTTTTCATAATAAATTGATTGCACTTTAAAGTAA |
| 7411_88 | A/T | 1 | CTGCAGTAAATTTTAAAAAAATGTAAATAAATTATCAGTAGCTTTAACACTTGATGGCAAAACGTGTTTAGGAAATACAATAATCGA[A/T]GGAAAGGGGCGTTTGGATCATGGATAGTGGGAGAGGATAAAAATTCCATTTCCATTTGTAACGTTGAAACGCGTTTTTCTTATGTTTGGAGTTATATTGATTATAAAAATGT |
| 788_100 | C/G | 1 | CTGCAGACTTGAGGTGTGGAGGGATTTTGTTAAGCAGTTTTGTGAATTTCATAATAAGGCTAGCCACTAATTCATTTTATTGCTTTTAAATGTTTGTGA[C/G]TTCATCTAGAACAATTTTTTGGGTCTCTGTGATCTCCATATTGTTCAACAAACATTCTTCTGAATTCAAGCCAGAAGCTGATTAGGGCATTTGGCAGATT |
| 8375_147 | A/T | 2 | CTGCAGAAAGAAAATGAAGTTGGAAATGGAATATATATAATTAGATTATAATTGGGTTCTAAAATATCTTTGGATGACATACTTTCGAACCTAGAAGAGCTGGATTTATCCAAGGTGGGCCAGTCTGATTGGAGATCCTTGAATTG[A/T]GCTTTGTAACCTCACTGGTCACCTTAGTCCAACACAGGATAATAGAGCACCTG |
| 9164_153 | G/A | 1 | CTGCAGATCAAGCATATAAATTTGAAAGAAAGGTAGTACAATGGATCCTATTTTCTTAACACTCTCAAGTCTGCTTTCTTTGGATAGACATCCTGTTTCTCGGTCCTCGCCAAAACATTCTCATCTCACCTTTATTAAATATCATTACAAAA[G/A]GAGGCGGCCTTTTACAAATAAGATGACTTTACTGTGAATGATGTCTC |
| 9258_25 | A/G | 2 | CTGCAGCATCTTGTGGTCCTCAAG[A/G]TCAATGAGGCAAATTGTGCACTCGAAGCCATTGAGCCCCTCCATAGATGTTCTTCATCAAGTTCCATCAAGGAGCTCATAAATCGCACCAATGATCAATAGTAATATATTCCTCTTCTGCTTTGATATTTGCCAAATCAAGAAGAGGTGCAATGATACAAGAGCATTATATTATG |
| 9590_145 | G/A | 1 | CTGCAGCACTTCCTTTCTGTTCGAAATTGAAAAAATGAATTATGGGTGAAGCTGCTTATTTTATTCCGAGTTTACCCTTTCAATAACCATTCGAAGGAAATCAATTAAGTCAAAACAGGTTGTAAATTTTGCAGCTAGAGCTGC[G/A]CAAAGAACTAGCTAGTAATAAGAAATAAAGCTTGTAGAAGAAAACAGACTGAGCT |
|  | 34 |  |  |
|  | 39 |  |  |

Appendix 2: Primer details for all 6 SSR loci used in this study. All forward primers had the M13 sequence added on the 5’ end of primer. JC035 was first described by (Michalczyk et al., 2006). NB: Loci 576s and 9104s were removed from final analyses for being null alleles

| Loci | Repeat motif | Repeat length | Amplicon size | Forward Primer Sequence | Reverse Primer Sequence |
| --- | --- | --- | --- | --- | --- |
| 8921s | (TACA) | 7 | 248 | TTCACGCGAAACTCTCCAG | AATGGCTTAACTGCCTCTCG |
| 8480s | (TATG) | 12 | 187 | CCTGCAATCGATGTCTTCCG | TGCCAAATAGTAACACAGATGGG |
| 576s | (ATGT) | 7 | 238 | GTAGACTCACTCTCCTCGAATG | TCCAGCAATTGATCACAAGGG |
| 9104s | (ATGT) | 8 | 152 | GTGTTGCTCCCATCTAGCAG | CCTTACACCACTGGATGCTC |
| 2346s | (TACA) | 7 | 229 | AGGTCCCACACCACTTATCC | CACATGGTCTTCCTTGGAGTTG |
| JC035 | (CA) | 20 | 155 | tgtgtttattctccccatct | cccccagttattctaaacatt |

Appendix 3: IDs (Populations), whether an individual was removed, and genotypes of all individuals who shared genotypes in the SNP dataset. Pairs of numbers are biallelic loci separated by dashes, numbers signify base pair: 1= A, 2=T, 3=G, 4=C. NB: only in those cases were individuals who shared a genotype and were from the same population was one of the clonal individuals removed

| Individual 1 | Individual 2 | Removed Individual | Genotype |
| --- | --- | --- | --- |
| 70 (CD) | 86 (CD) | 70 | 12/21/33/42/44/31/24/11/11/11/11/11/22/11/11/43/33/11/22/42/21/11/22/44/42/44/33/22/44/33/33/41/33/31/33/31/42/22/11/31/11 |
| 71 (CD) | 83 (CD) | 71 | 22/21/31/22/44/11/24/11/11/11/44/11/21/11/31/43/33/13/22/44/11/11/22/44/44/44/33/33/44/11/11/11/33/31/33/33/42/11/11/33/11 |
| 213 (Nana) | 214 (Nana) | 213 | 22/22/31/22/44/11/24/13/11/44/11/11/22/11/31/43/33/13/22/44/11/31/22/33/22/44/33/32/44/33/11/41/31/31/33/33/44/11/11/33/11 |
| 214 (Nana) | 216 (Nana) | 214 | 22/22/31/22/44/11/24/13/11/44/11/11/22/11/31/43/33/13/22/44/11/31/22/33/22/44/33/32/44/33/11/41/31/31/33/33/44/11/11/33/11 |
| 157 (Hemi) | 158 (Hemi) | 157 | 12/22/31/22/44/33/24/11/41/44/41/11/21/33/31/44/33/13/22/42/11/31/22/44/22/44/33/33/22/33/11/44/33/33/33/33/44/11/11/33/31 |
| 158 (Hemi) | 159 (Hemi) | 158 | 12/22/31/22/44/33/24/11/41/44/41/11/21/33/31/44/33/13/22/42/11/31/22/44/22/44/33/33/22/33/11/44/33/33/33/33/44/11/11/33/31 |
| 162 (Hemi) | 167 (Hemi) | 162 | 22/21/33/22/44/33/22/11/11/41/41/11/21/33/11/44/33/13/22/42/11/31/22/44/22/44/33/33/22/33/11/44/33/33/33/33/44/11/11/33/31 |

Appendix 4: IDs (Populations), whether an individual was removed, and genotypes of all individuals who shared genotypes in the SSR dataset. Numbers are the number of base pairs at each loci, with alleles separated with dash. NB: only in those cases were individuals who shared a genotype and were from the same population was one of the clonal individuals removed

| Individual 1 | Individual 2 | Removed individual | Locus 1 | Locus 2 | Locus 3 | Locus 4 |
| --- | --- | --- | --- | --- | --- | --- |
| 22 (BG) | 224 (NANA) | n/a | 250/250 | 212/196 | 230/222 | 250/250 |
| 32 (BT) | 280 (BF) | n/a | 258/258 | 200/196 | 224/224 | 254/250 |
| 38 (BT) | 228 (TF) | n/a | 258/250 | 212/200 | 224/224 | 250/250 |
| 45 (BF) | 15 (BG) | n/a | 250/250 | 200/196 | 224/224 | 250/250 |
| 54 (BH) | 27 (BG) | n/a | 250/250 | 208/200 | 224/222 | 250/250 |
| 61 (CD) | 79 (CD) | 61 | 250/250 | 208/200 | 230/226 | 250/250 |
| 62 (CD) | 84 (CD) | 62 | 226/258 | 200/196 | 230/224 | 250/250 |
| 66 (CD) | 83 (CD) | 83 | 258/254 | 216/196 | 226/226 | 254/250 |
| 67 (CD) | 85 (CD) | 67 | 258/250 | 200/196 | 222/222 | 250/250 |
| 96 (CD) | 210 (LM) | n/a | 258/250 | 200/196 | 226/224 | 250/250 |
| 150 (FK) | 181 (GA) | n/a | 250/250 | 196/196 | 226/224 | 250/234 |
| 187 (HEMI) | 193 (HEMI) | 187 | 250/250 | 232/196 | 224/224 | 238/238 |
| 189(HEMI) | 192 (HEMI) | 189 | 250/250 | 232/232 | 224/224 | 254/238 |
| 210 (LM) | 211 (LM) | 211 | 258/250 | 200/196 | 226/224 | 250/250 |
| 211 (LM) | 313 (WW) | n/a | 258/250 | 200/196 | 226/224 | 250/250 |
| 222 (LM) | 226 (LM) | 222 | 258/250 | 200/200 | 224/224 | 250/250 |
| 243 (NANA) | 245 (NANA) | 243 | 250/250 | 216/196 | 230/224 | 250/250 |
| 258 (PD) | 261 (PD) | 258 | 258/258 | 196/196 | 226/222 | 250/250 |
